# Nuclear stress bodies enable a germline-specific transcriptional stress response in *Drosophila*

**DOI:** 10.64898/2026.08.24.746661

**Authors:** Dhanashree Lakhe, Lorenzo Gallicchio, Juan Sebastian Ochoa Gutierrez, Joshua Düring, Demir Erensoy, Franziska Brändle, Anna Sintsova, J. Matthew Franklin, Madhav Jagannathan

## Abstract

The germline ensures the continuity of genetic information across generations, but how this immortal lineage functions under stress conditions remains incompletely understood. Here, we identify that heat shock factor (Hsf), a conserved master regulator of the stress response, drives the expression of transposable elements (TEs), in addition to molecular chaperones upon heat shock in *Drosophila* gonads. In germ cells, this potential intra-genomic conflict is countered by the formation of nuclear stress bodies (nSBs) at non-coding satellite DNA repeats. Using chemical and genetic perturbations, we demonstrate that nSBs are both necessary and sufficient to delay Hsf-dependent transcription. Notably, this nSB-mediated delay, in tandem with the piRNA pathway, allows germ cells to selectively express molecular chaperones, but not transposable elements, upon heat shock. Overall, we propose that this unique transcriptional stress response preserves germline function and evolutionary fitness, especially in natural populations routinely exposed to environmental stress.

## Introduction

Reproduction in many multicellular organisms requires a specialized cellular lineage known as the germline^1^. Since germ cells are the sole carriers of genetic information across generations, the maintenance of germline genome integrity is imperative to avoid the inheritance of deleterious mutations^2^. A number of cellular features ensure the preservation of genome integrity in the germline^1,3^. For example, relatively low rates of replication and transcription in germ cells ensure that mutations associated with these processes are minimized^1^. In addition, germ cells possess a hyperactive response to DNA damage, often triggering cell death rather than risking the transmission of mutations to the offspring^4–8^. Finally, germ cells deploy a dedicated small RNA pathway known as the piRNA pathway to counteract the expression of mobile and genotoxic transposable elements (TEs)^9,10^. When germline genome integrity is compromised, individuals can exhibit reduced fertility and their offspring can experience pathological outcomes.

Beyond specialized features that maintain genome integrity, the germline also behaves differently from the soma in other cellular processes, including the environmental stress response^11^. Organisms routinely encounter stressors, such as heat shock, osmotic shock, nutrient starvation, in their natural habitats. These diverse stressors trigger a rapid and specific transcriptional response involving the expression of molecular chaperones via the conserved heat shock factor (Hsf) family of transcription factors^12,13^. This stress response, often called the heat shock response, serves as a first line of defense against protein misfolding and aggregation and is required for survival during stress^12,13^. However, certain aspects of the canonical stress response are known to impair germline function. For example, constitutively active Hsf induces germ cell apoptosis and spermatogenesis failure in mammals^14–16^. In *Drosophila*, stress-induced overexpression of Hsp70 chaperones induces the degradation of proteins that function in TE repression^17^ and reduces germ cell viability^18^. Intriguingly, stress also triggers TE expression across several species^19^, including plants^20–23^, and animals^17,24–29^. How germ cells balance undesirable effects of the stress response while maintaining their genome integrity and reproductive output is not fully understood.

In this study, we identify that germ cells exhibit a distinctive transcriptional response to stress in *Drosophila*. We find that Hsf, the conserved transcription factor that normally drives chaperone expression under stress conditions, also stimulates the expression of chaperone-proximal TE insertions, setting off an intra-genomic conflict that could potentially harm germline genome integrity. In germ cells, we show that this threat is mitigated by the formation of dynamic Hsf-containing nuclear stress bodies (nSBs) at non-coding satellite DNA repeats. We demonstrate that these nuclear foci are both necessary and sufficient to delay the immediate transcriptional response to heat shock, enabling selective chaperone expression upon nSB disassembly. In the absence of nSBs, germ cells express both chaperones and TEs immediately upon heat shock and exhibit elevated cell death at later time points. Notably, modulating satellite DNA abundance and accessibility modifies nSB formation and function, highlighting a previously unappreciated role for these repeats in the germline stress response. Overall, our work reveals a germ cell-specific mechanism that maintains robust gametogenesis under stress conditions.

## Results

### A germline-specific transcriptional response to stress

We set out to characterize the transcriptional response to stress in *Drosophila* germ cells. To do so, we used heat shock as a stressor, subjecting adult flies to a one-hour heat shock at 37°C. After allowing flies to recover for one hour at 25°C, we dissected testes, extracted RNA and performed RNA sequencing (Fig. 1A). To minimize transcriptional variability arising from the various stages of male germline development, we used flies with testes enriched for mitotically proliferating germ cells, hereafter spermatogonia, due to a mutation in the *bag of marbles* (*bam*) gene^30^. We observed that 199 genes were upregulated upon heat shock (log_2_FC >1, p_adj_<0.05), including several chaperones such as *Heat shock protein 26* (*Hsp26*), *Hsp68* and all six *Hsp70* paralogues in the *Drosophila* genome (Fig. 1B, cyan circles). We also observed heat shock-induced expression of two TE families, specifically the internal sequence of the *copia2* retrotransposon (*copia2_I*) and the long terminal repeat (LTR) regions of the *invader1* retrotransposon (*invader1_LTR*), in spermatogonia-enriched testes (Fig. 1B, magenta circles). Out of the 32 insertions of *copia2* (also known as *Dm88*) and 26 insertions of *invader1* across the *Drosophila* euchromatin, only one *invader1* element is full-length and presumably capable of mobilization^31^. Notably, both TE families are also expressed upon heat shock in female gonads^27^. Thus, stress induces the expression of several chaperones and two TE families in *Drosophila* gonads.

**Figure 1.**
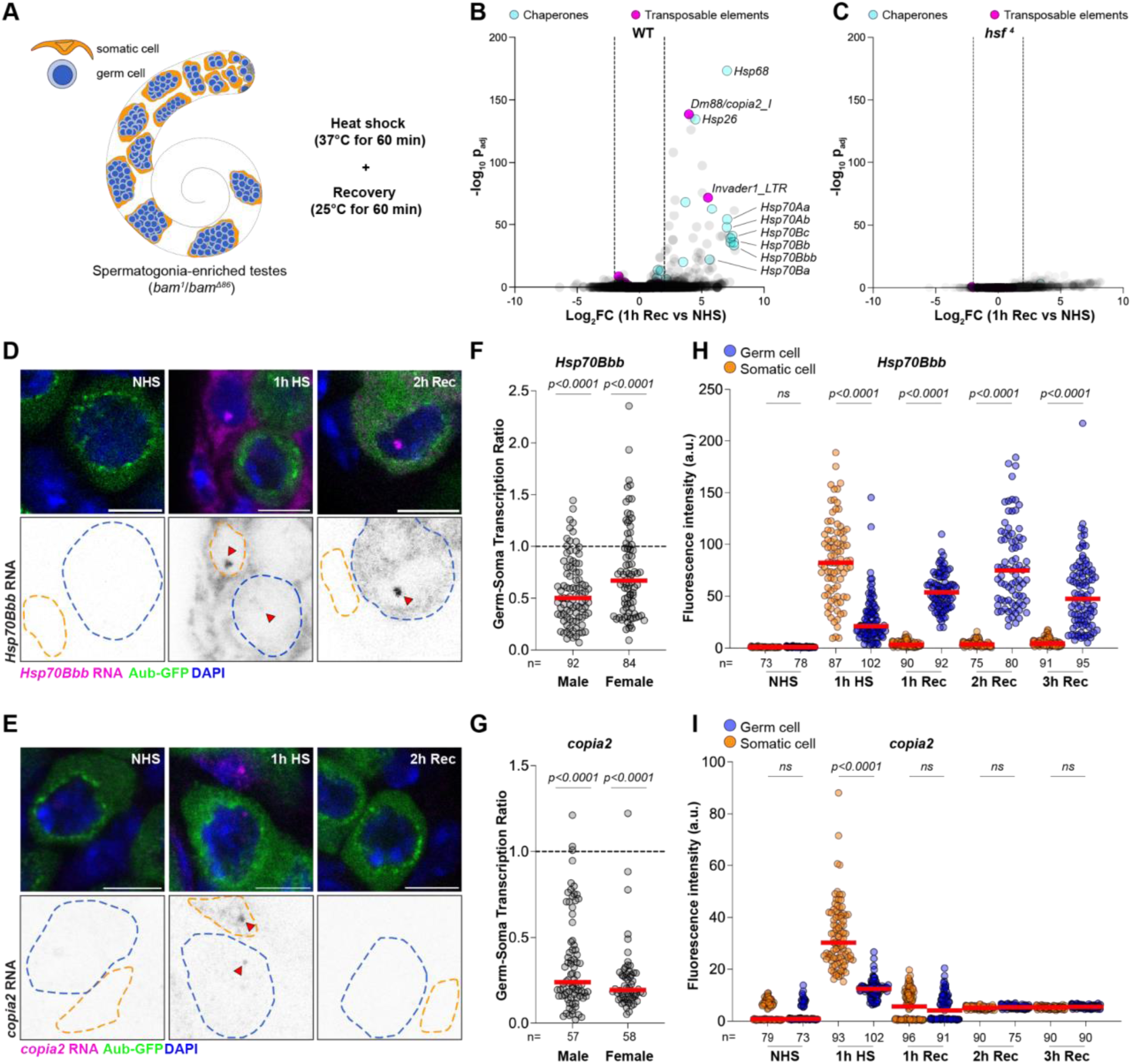
The germline exhibits a delayed and selective transcriptional stress response. (A) A schematic of the heat shock regimen before RNA extraction from spermatogonia-enriched (*bam*^1^*/bam^Δ^*^86^) testes. (B,C) Volcano plots of the transcriptome from *WT* (B) or *hsf* ^4^ (C) spermatogonia-enriched testes before and after heat shock. Cyan circles represent chaperones and pink circles represent transposable element families. (D, E) RNA FISH against *Hsp70Bbb* (D) or *copia2* (E) transcripts (magenta) at the indicated time points in Aub-GFP (green) testes stained with DAPI (blue). Blue dashed lines demarcate germ cells while orange dashed lines demarcate somatic cell nuclei. Arrowheads indicate nascent transcriptional loci. Scale bar:5µm (F,G) Quantification of the germ-soma transcription ratio in mitotically proliferating male and female germ cells for *Hsp70Bbb* (F) and *copia2* (G). Red lines indicate median values. *p*-values obtained from a one sample *t*-test against a hypothetical value of 1. (H,I) Quantification of cytoplasmic *Hsp70Bbb* (H) and nucleoplasmic *copia2* (I) transcripts based on RNA FISH fluorescent intensities in somatic (orange) and germ (blue) cells from testes at the indicated time points. Red lines indicate median values. *p-*values were obtained from a two-way ANOVA. ns indicates *p*>0.05.

We next sought to determine the molecular factors responsible for the stress-induced expression of chaperones and TEs. Across eukaryotes, the transcriptional response to stress involves evolutionarily conserved transcription factors known as Heat shock factors^12,13^. The *Drosophila* genome contains a single Heat shock factor (Hsf) gene, which is essential for development^32^. However, a previously described temperature-sensitive allele called *hsf* ^4^, which contains a mutation in the DNA-binding domain, renders Hsf inactive at and above 29°C^32^, thereby allowing conditional abrogation of Hsf function. In spermatogonia-enriched *hsf* ^4^ testes, we did not observe upregulation of any chaperones upon heat shock (Fig. 1C), consistent with the known role of Hsf. Unexpectedly, loss of Hsf function also abolished stress-induced expression of *copia2* and *invader1*, indicating that the same transcription factor, i.e. Hsf, is responsible for the expression of both chaperones and TEs upon heat shock.

Spermatogonia-enriched *Drosophila* testes contain both germline and supporting somatic cells and our bulk RNAseq dataset does not reveal whether the observed stress-induced expression of TEs and chaperones occurs in the germline, soma or in both populations. Therefore, we used single molecule RNA FISH to label stress-induced transcripts from *Hsp70Bbb,* a representative chaperone, and the *copia2* TE in order to determine their spatio-temporal expression patterns upon heat shock. In an *Aubergine-GFP* (*Aub-GFP*) strain that specifically marks the germ cell cytoplasm, we found that neither *Hsp70Bbb* nor *copia2* were transcribed in the absence of heat shock (Fig. 1D, E, NHS). Immediately upon heat shock (1h HS), we observed that spermatogonia had a weaker nascent transcription signal compared to the somatic cells in the same tissue (Fig. 1D, E, arrowheads point to nascent transcription loci, Fig. S1A, B). To minimize variability arising from potential differences in imaging conditions and sample preparation, we developed a metric called the germ-soma transcription ratio (Fig. S1C), where we divide the total fluorescent intensity of the nascent locus in a germ cell by that of a nascent locus in a somatic cell in the same or adjacent imaging plane (see Methods for details). A germ-soma transcription ratio below 1 indicates that the germ cells have a weaker transcriptional response to stress while values above 1 indicate the opposite. Spermatogonia exhibited a median germ-soma transcription ratio of 0.503 for *Hsp70Bbb* and 0.349 for *copia2* (Fig. 1F, G), both significantly below 1, indicating a weaker Hsf-dependent transcriptional response in comparison to their neighbouring somatic cells. Similarly, nascent chaperone and TE transcription was diminished in mitotically proliferating female early germ cells in comparison to the surrounding somatic cells (Fig. 1F, G) immediately upon heat shock. These data suggest that the Hsf-dependent and stress-induced expression of chaperones and TEs detected in our RNAseq data primarily originate from somatic cells.

To assess the transcriptional response to heat shock at later time points when nascent loci are absent, we also quantified the expression of *Hsp70Bbb* (cytoplasmic) and expression of *copia2* (nucleoplasmic) by RNA FISH during the recovery from heat shock. To unambiguously determine the boundaries of the germ cells from the somatic cells that engulf them, we labelled the somatic cell cytoplasm using cell type-specific expression of a fluorescent ribosomal protein (*c587-Gal4* > *UASt-GFP-RpL10Ab*) (Fig. S1D). As before, spermatogonia in this genetic background exhibited a median germ-soma transcription ratio below 1 (Fig. S1E). Immediately upon heat shock (1h HS), somatic cells displayed prominent cytoplasmic *Hsp70Bbb* transcripts (Fig. 1D, H Fig. S1F). At the same time point, *Hsp70Bbb* transcripts were only weakly detected in the germ cell cytoplasm, but gradually increased in expression during recovery, peaking at 2h post heat shock (2h Rec) (Fig. 1D, H, Fig. S1F). For *copia2*, we detected high levels of nucleoplasmic transcripts in somatic cells immediately upon heat shock (Fig. 1E, I), that did not persist during recovery. Interestingly, and in marked contrast to *Hsp70Bbb*, we did not observe nucleoplasmic *copia2* expression in germ cells during recovery from heat shock (Fig. 1E, I). In female germ cells, we observed a similar delay in *Hsp70Bbb* expression and an absence of *copia2* expression following heat shock (Fig. S1G-J).

Overall, our data reveal that early germ cells have a distinct transcriptional response to stress. Immediately upon heat shock, they exhibit diminished chaperone and TE expression in comparison to their surrounding somatic cells. Interestingly, *Hsp70Bbb* expression builds in germ cells during recovery, suggesting a delayed transcriptional response to stress. Moreover, this delayed transcriptional response appears to be selective, eventually leading to the expression of the *Hsp70Bbb* chaperone in germ cells, but not the *copia2* TE.

### Chaperone-embedded TEs are specifically expressed upon stress

We first wanted to understand why heat shock only induced expression of two (*copia2* and *invader1*) out of the 135 TE families in the *Drosophila* genome^31,33^. We noticed that 17 *copia2* and 15 *invader1* insertions were embedded within a locus flanked by four *Hsp70* paralogues on Chr. 3R (Fig 2A, B, *Hsp70B* locus, yellow and red insertions, Fig. S2A). Although short reads from NGS datasets can be challenging to assign to specific TE insertions due to multimapping, we noted that 11 individual TE insertions were upregulated upon heat shock in our transcriptomic analysis (Fig. S2B). Of these, 7 TE insertions were embedded in the *Hsp70B* locus (Fig. S2B), with their proximity to unique *Hsp70* coding sequences presumably aiding their identification. This locus also contained several LTR fragments of *invader1* (Fig. S2A), which likely explains the detection of the *invader1 LTR* transcripts in our RNAseq data (Fig. 1B).

**Figure 2.**
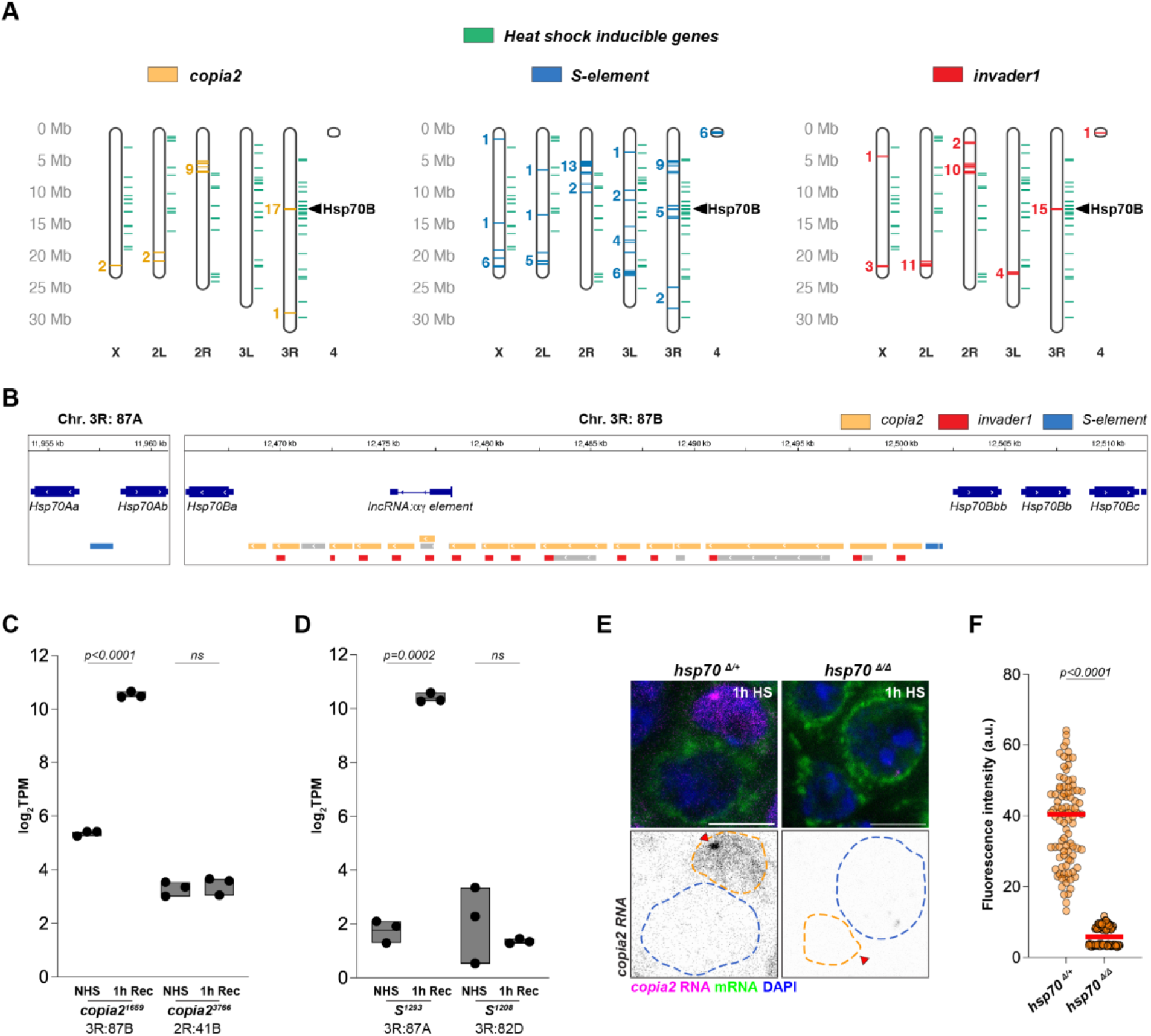
Chaperone-embedded TEs are specifically expressed upon stress. (A) Genomic locations of *copia2* (yellow), *S-element* (blue) and *invader1* (red) insertions. Green lines indicate the location of heat shock-induced genes from our RNA-seq dataset. (B) A representation of the *Hsp70A* and *Hps70B* locus at cytological bands 87A and 87C on Chr. 3R. *Hsp70* genes are depicted in dark blue, *copia2* insertions in yellow, *S-element* insertions in blue and *invader1* insertions in red. (C,D) Normalized read counts of *copia2* (C) and *S-element* (D) insertions within and outside the *Hsp70* gene loci from the RNAseq dataset. *p-*values were obtained from a Welch’s t-test. ns indicates *p*>0.05. (E) RNA FISH against *copia2* transcripts (magenta) and mRNA (green) in testes carrying a deletion of the *Hsp70* gene cluster (*hsp70^Δ/Δ^*) compared to a heterozygous control (*hsp70^Δ/+^*). Blue dashed lines demarcate germ cells while orange dashed lines demarcate somatic cell nuclei. Scale bars:5µm (F) Quantification of nucleoplasmic *copia2* expression in somatic cells from (E). *p*-value was obtained from a Welch’s t-test.

Moreover, we found that a *copia2* insertion from within the *Hsp70B* locus was highly expressed upon heat shock, unlike a representative insertion present elsewhere in the genome (Fig. 2C). This suggested a potential relationship between the *Hsp70*-proximal position of the TE and its stress-induced expression. If true, other TE insertions in the *Hsp70B* locus would also be expected to be expressed upon heat shock. Indeed, this locus also contains two insertions of the *S-element*, a terminal inverted repeat (TIR)-containing DNA transposon (Fig. 2A, C). In addition, a single S-element fragment is embedded between the other two *Hsp70* paralogues, *Hsp70Aa* and *Hsp70Ab* (*Hsp70A* locus, Fig. 2B). We observed from our RNAseq data that all three S-elements were upregulated upon heat shock (Fig. 2D, Fig. S2B), in comparison to S-element insertions at other genomic loci (Fig. 2D).

The above data suggested that the Hsp70-proximal TEs are the major source of the nucleoplasmic *copia2* signal observed in somatic cells upon heat shock. To test this, we used a strain containing a precise deletion of the *Hsp70A* and *Hsp70B* gene clusters including the embedded TEs (*hsp70^Δ/Δ^*). In flies homozygous for this deletion, we did not detect heat shock-induced *copia2* expression, in comparison to the heterozygous control (Fig 2E, F). Altogether, these data suggest that it is the presence within a Hsf regulated locus, rather than underlying regulatory sequences, that is the primary determinant of stress-induced TE expression.

### Hsf-containing nuclear stress bodies are specifically formed in germ cells and delay chaperone expression

We next set out to elucidate the mechanistic basis underlying the delayed or diminished expression of *Hsp70Bbb* and *copia2* in germ cells following heat shock. Hsf-dependent transcription under stress conditions is regulated by several mechanisms, including nuclear localization, trimerization, chaperone titration, post-translational modifications and the chromatin state at target loci^12,13,34–41^. In addition, micron-sized Hsf foci known as nuclear stress bodies (nSBs) have been demonstrated to impair chaperone expression, primarily in cancer cells^42–46^. In the absence of stress, we found that Hsf was uniformly localized within somatic and germ cell nuclei in testes and ovaries (Fig. 3A, B). However, upon heat shock, we observed micron-sized Hsf foci specifically in male and female germ cells, while Hsf localization remained unchanged in somatic cells (Fig. 3A-C). The number and size of these Hsf foci was consistent in male and female germ cells and different stages of spermatogonial development (Fig. S3A-D). Similar foci were also observed in undifferentiated germ cells in larval spermatogonia (Fig. S3E) and germ cells from stage 15-16 embryos (Fig. S3F) upon heat shock, suggesting that germ cells across all stages of development form these structures. In contrast, stress-induced Hsf foci were not observed in all larval somatic tissues tested (Fig. S3G-K). These foci resemble previously reported Hsf-containing nuclear stress bodies (nSBs), which form on non-coding tandem repeats known as satellite DNA and have thus far only been observed in somatic, and typically transformed, primate cells^42,47,48^. In addition to Hsf, primate nSBs are enriched for several transcriptional regulators, including RNA polymerase II^42,49,50^, and their formation is associated with diminished Hsf activity at chaperone loci^43^.

**Figure 3.**
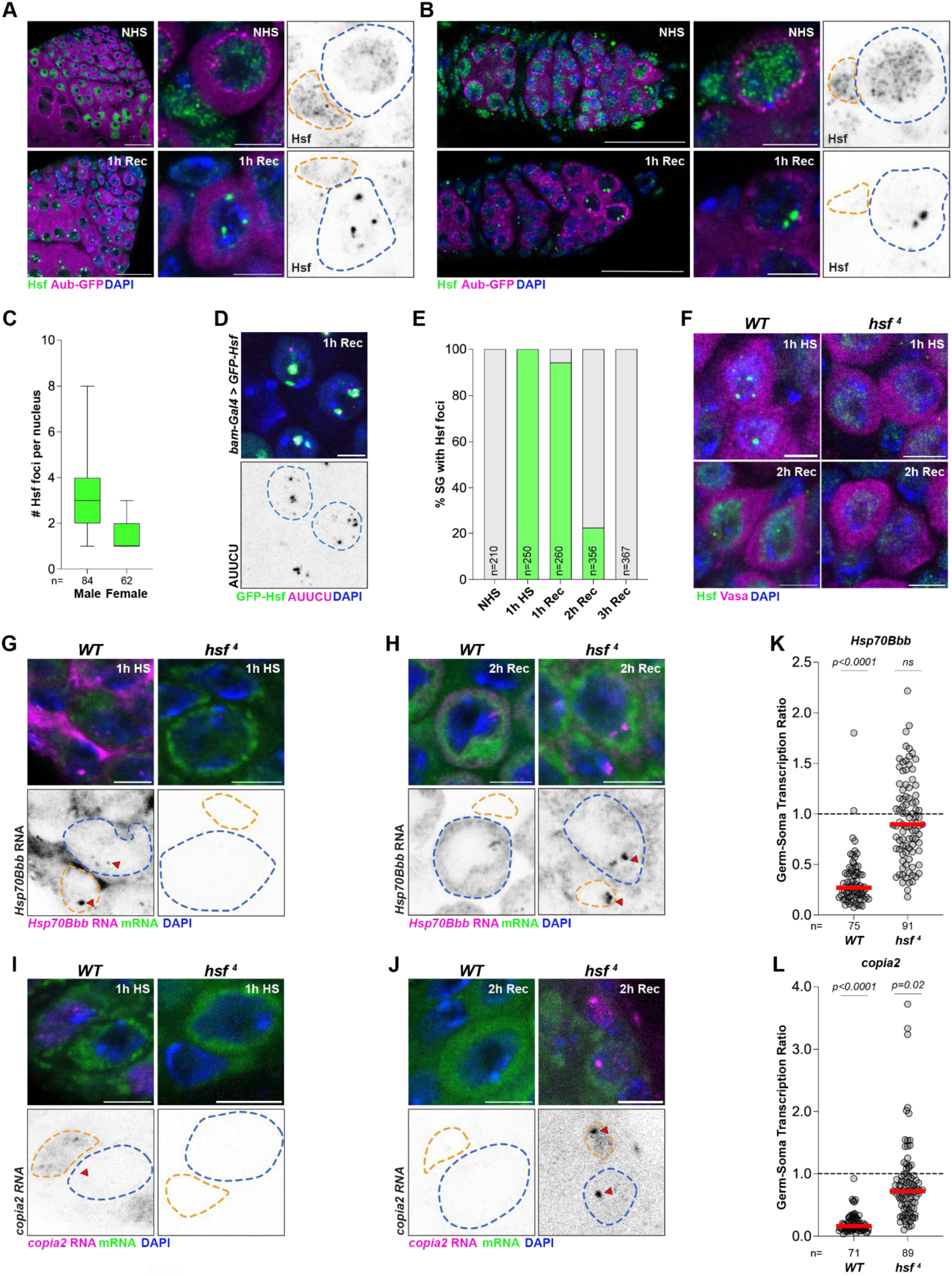
Hsf-containing nuclear stress bodies are specifically formed in germ cells. (A,B) Apical tip of the testes (A) and germarium of the ovaries (B) at the indicated time points stained for Hsf (green) and DAPI (blue) in an Aub-GFP (magenta) background. Scale bars:25µm and 5µm for insets. (C) Quantification of number of Hsf foci per nucleus in male and female mitotically proliferating germ cells. (D) RNA FISH detecting ATTCT transcripts in male germ cells expressing GFP-Hsf under the control of *bam-Gal4*. Scale bar:5µm (E) Quantification of the percentage of spermatogonia with Hsf foci at the indicated time points in a *c587-Gal4*>*UAS-mCD4-tdTomato* background. (F) Testes from *Oregon R* and *hsf* ^4^ stained for Hsf (green), Vasa (magenta) and DAPI (blue) at the indicated time points. Scale bars:5µm. (G-J) RNA FISH against *Hsp70Bbb* (G, H) and *copia2* (I, J) transcripts in male germ cells from the indicated genotypes at 1h HS (G, I) and 2h Rec (H, J). Scale bars:5µm (K,l) Quantification of the germ-soma transcription ratio for *Hsp70Bbb* (K) and *copia2* (L) in WT (1h HS) and *hsf* ^4^ (2h Rec) testes. Red lines indicate median values. *p*-values obtained from a one-sample *t*-test against a hypothetical value of 1.

In human cells, nSBs are formed at the HSat3 satellite DNA^42^, whose GGAAT repeats resemble the conserved NGAAN containing Hsf binding motif at chaperone promoters^35,51–53^. As a result, Hsf and its associated transcriptional regulators drive the transcription of Hsat3, with these transcripts remaining localized at nSBs^48,49^. Not only are these satellite RNAs a hallmark feature of nuclear stress bodies, their sequence also reveals the nSB-nucleating satellite DNA repeat. To determine whether the germ cell-specific Hsf foci were associated with satellite DNA transcription, we reanalysed our RNAseq dataset using an assembly-independent k-mer detection pipeline called k-seek^54^, which quantifies 2-25bp tandem repeats present in NGS reads. We observed that the AUUCU k-mer was highly enriched upon heat shock in WT testes, but not in the *hsf*^4^ background (Fig. S3L), suggesting that Hsf is required for its expression. Notably, AUUCU corresponds to an approximately 1Mb satellite DNA repeat, AGAAT/ATTCT, which is primarily present on Chr. 2 and Chr. Y^55,56^. Moreover, we detected AUUCU transcripts that were localized to Hsf foci in germ cells using RNA FISH (Fig. 3D). Collectively, our data indicate that the germ cell-specific Hsf foci are formed at the AGAAT satellite DNA repeat upon heat shock and we refer to them hereafter as *Drosophila* nuclear stress bodies (nSBs).

nSB formation in human cells is associated with diminished Hsf activity^43^, leading us to speculate that *Drosophila* nSBs may be responsible for the transcriptional delay observed in germ cells. Consistently, *Drosophila* nSBs, which form within minutes of exposure to heat shock (Fig. S3M), are disassembled approximately two hours post stress (Fig. 3E), which coincides with *Hsp70Bbb* expression in germ cells (Fig. 1H). Unlike in *WT* (*Oregon R*) germ cells, nSBs were not observed in *hsf* ^4^ mutant germ cells, either during or post heat shock (Fig. 3F), likely due to impaired DNA-binding at the non-permissive temperature. However, we observed *Hsp70Bbb* and *copia2* expression in *hsf* ^4^ germ and somatic cells around 1-2h post heat shock, likely due to the restoration of Hsf function at the permissive temperature (Fig. 3G-J). Interestingly, the germ-soma transcription ratio in *hsf* ^4^ testes at 2h post heat shock was close to 1, for both *Hsp70Bbb* and *copia2* (Fig. 3K, L), suggesting that germ cells and somatic cells have a similar Hsf-dependent transcriptional response to heat shock in the absence of nSBs. These data support the idea that nSB formation in germ cells delays the transcriptional response to stress in comparison to the neighbouring soma.

### Hsf-containing nSBs are necessary and sufficient for the germ cell-specific transcriptional response

We next set out to more directly manipulate nSBs to test their role in stress-induced *Hsp70Bbb* and *copia2* expression. First, we blocked nSB formation using *ex vivo* treatment with a known chemical inhibitor, mitoxantrone^43^ (see Methods). We observed that mitoxantrone blocked nSB formation in germ cells (Fig. 4A, B), which corresponded to a significant increase in the germ-soma transcription ratio for both *Hsp70Bbb* (Fig.4C, D) and *copia2* (Fig. 4E, F) in comparison to the mock-treated control. These data indicate that nSBs are necessary for diminishing chaperone and TE expression in germ cells, immediately upon heat shock.

**Figure 4.**
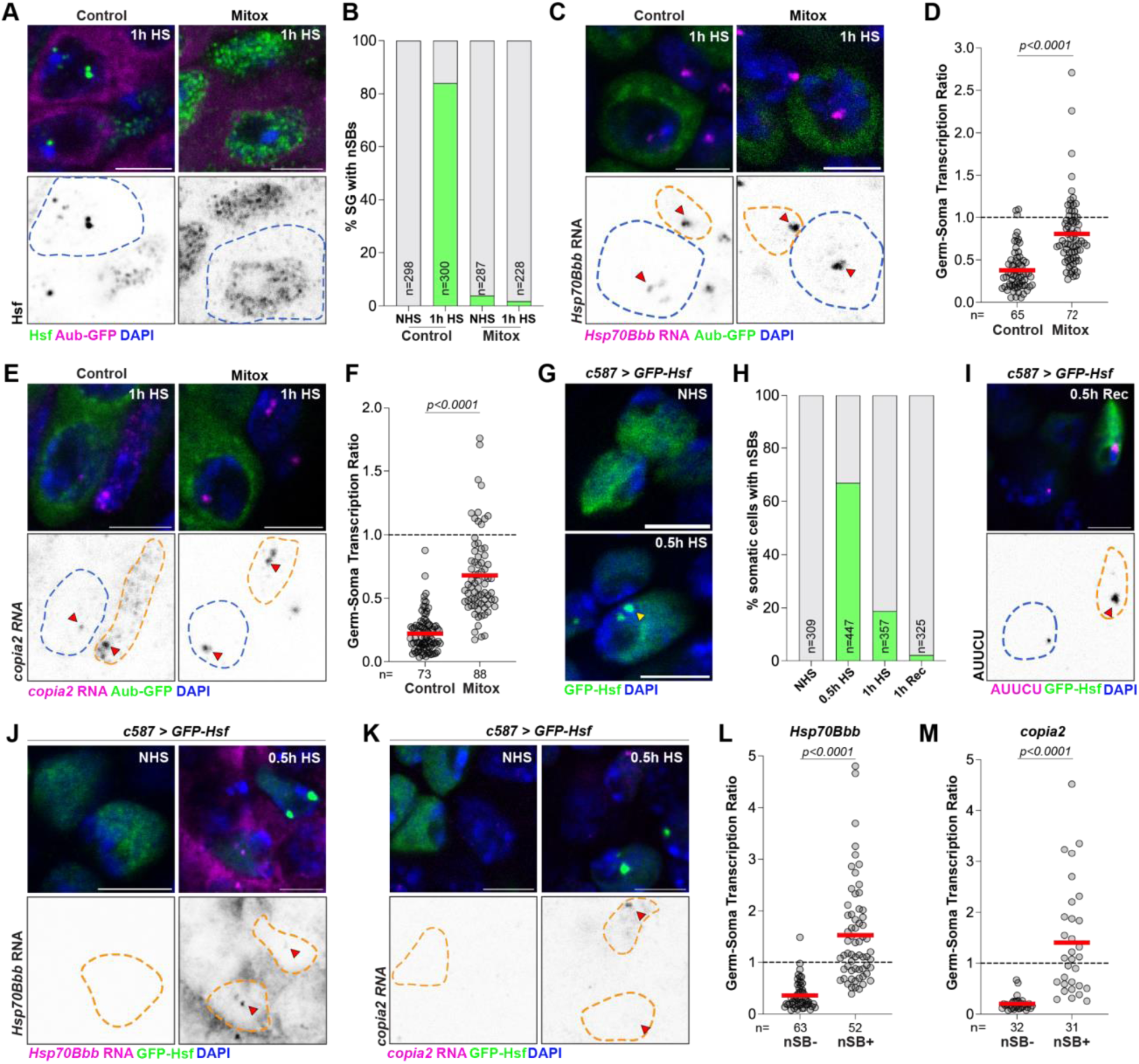
Hsf-containing nSBs are necessary and sufficient for the germ cell-specific transcriptional response to stress. (A) Spermatogonia from Aub-GFP (magenta) testes stained for Hsf (green) and DAPI (blue) after ex vivo treatment with ddH2O (Control) or 20 µm Mitoxantrone (Mitox) followed by a 1h HS. Blue dashed lines demarcate germ cells. Scale bars:5µm. (B) Quantification of percentage of spermatogonia with nSBs in the indicated conditions. (C) RNA FISH against *Hsp70Bbb* transcripts (magenta) in male germ cells expressing Aub-GFP (green) and stained for DAPI (blue) in the indicated conditions at 1h HS. Blue dashed lines demarcate germ cells while orange dashed lines demarcate somatic cell nuclei. Arrowheads indicate nascent transcriptional loci. Scale bars:5µm (D) Quantification of the germ-soma transcription ratio for *Hsp70Bbb* from (C) at 1h HS. Red lines indicate median values. *p*-value obtained from a Welch’s t-test. (E) RNA FISH against *copia2* transcripts (magenta) in male germ cells expressing Aub-GFP (green) and stained for DAPI (blue) in the indicated conditions at 1h HS. Arrowheads indicate nascent transcriptional loci. Scale bars:5µm (F) Quantification of the germ-soma transcription ratio for *copia2* from (E) at 1h HS. Red lines indicate median values. *p*-value obtained from a Welch’s t-test. (G) Somatic cysts cells from testes expressing *GFP-Hsf* (green) under the control of *c587-Gal4* and stained for DAPI (blue) at the indicated time points. Yellow arrowhead marks somatic nSBs. Scale bars:5µm. (H) Percentage of somatic cells from (G) that form nSBs at the indicated time points. (I) RNA FISH detecting ATTCT transcripts from (G) at 0.5h Rec. Scale bar:5µm. (J, K) RNA FISH against *Hsp70Bbb* (J) and *copia2* (K) transcripts (magenta) from testes stained for DAPI (blue) containing somatic cyst cells with and without Hsf-containing nSBs (green) at the indicated time points. Orange dashed lines demarcate somatic nuclei while arrowheads indicate nascent transcriptional loci. Scale bars:5µm (L, M) Quantification of the germ-soma transcription ratio for *Hsp70Bbb* (L) and *copia2* (M) from nSB-containing (nSB+) and nSB-lacking (nSB-) somatic cyst cells from (J, K) at 0.5h HS. Red lines indicate median values. *p*-values obtained from a Welch’s t-test.

Unexpectedly, we also found that Hsf overexpression (*c587-Gal4* > *GFP-Hsf*) led to transient nSB formation in a subset of somatic cyst cells (Fig. 4G, H). The majority of these somatic nSBs only persisted for 30-60 minutes post heat shock (Fig. 4G, H) and their formation was associated with AGAAT satellite DNA transcription (Fig. 4I). Strikingly, nSB-positive somatic cells exhibited higher germ-soma transcription ratio values for both *Hsp70Bbb* and *copia2* in comparison to neighbouring nSB-negative somatic cells (Fig. 4J-M). Altogether, we conclude that the formation of nSBs at satellite DNA repeats underlies the delayed transcriptional response to stress in germ cells.

### The piRNA pathway ensures silencing of chaperone-embedded TEs during the recovery from heat shock

During recovery from heat shock, the disassembly of Hsf-containing nSBs in germ cells coincided with the expression of *Hsp70Bbb* but not *copia2* in germ cells (Fig. 1H, I), and the basis for this selectivity remained unknown. Since TEs, including chaperone-proximal TE loci, are generally decorated with constitutive heterochromatin marks like H3K9me3^57,58^, we hypothesized that this difference in the chromatin state may underlie the selective expression by Hsf (Fig. 5A). In animals, a small RNA pathway known as the Piwi-interacting RNA (piRNA) pathway silences TE expression in the germline through heterochromatin formation at TE loci and transcript degradation in the cytosol^9,10^. In *Drosophila* germ cells, piRNA precursor transcription relies on the Heterochromatin Protein 1 (HP1) paralogue, Rhino, in complex with Deadlock and Cutoff (Fig. 5A)^58,59^. Following processing in the cytoplasm, mature piRNAs are loaded onto RNA-binding Argonaute proteins including Piwi (P-element induced wimpy testis)^9,10^. piRNA-bound Piwi enters the nucleus, locates TE loci through sequence complementarity and facilitates local H3K9me3 heterochromatin formation with the help of the SFiNX/PANDAS/PICTS complex (Fig. 5A)^60–64^. As such, loss of either Rhino or Panoramix (Panx), part of the SFiNX/PANDAS/PICTS complex, leads to loss of TE-associated heterochromatin^58,59,61,62^. To test whether TE-associated and piRNA-dependent heterochromatin was responsible for the selective expression of *Hsp70Bbb* during the recovery from heat shock, we used validated RNAi strains to deplete either Rhino or Panx. Immediately upon heat shock, nSB formation and *copia2* expression were quite similar between Rhino- and Panx-depleted germ cells and the mCherry RNAi control (Fig. S4A-E). During recovery (2h Rec), we observed that depletion of Rhino and Panx led to *copia2* expression in germ cells (Fig. 5B, C), with up to 60% of cells exhibiting nucleoplasmic signal (Fig. 5D). These data suggested that piRNA-mediated heterochromatin deposition at TE loci is crucial for maintaining *copia2* silencing post heat shock. We note, however, that the likely presence of H3K9me3 heterochromatin at TE loci in somatic cells (Fig. 1E, I), or in nSB-deficient germ cells (Fig. 4E, F), does not block Hsf-driven *copia2* expression, possibly due to impaired heterochromatin function at elevated temperatures.

**Figure 5.**
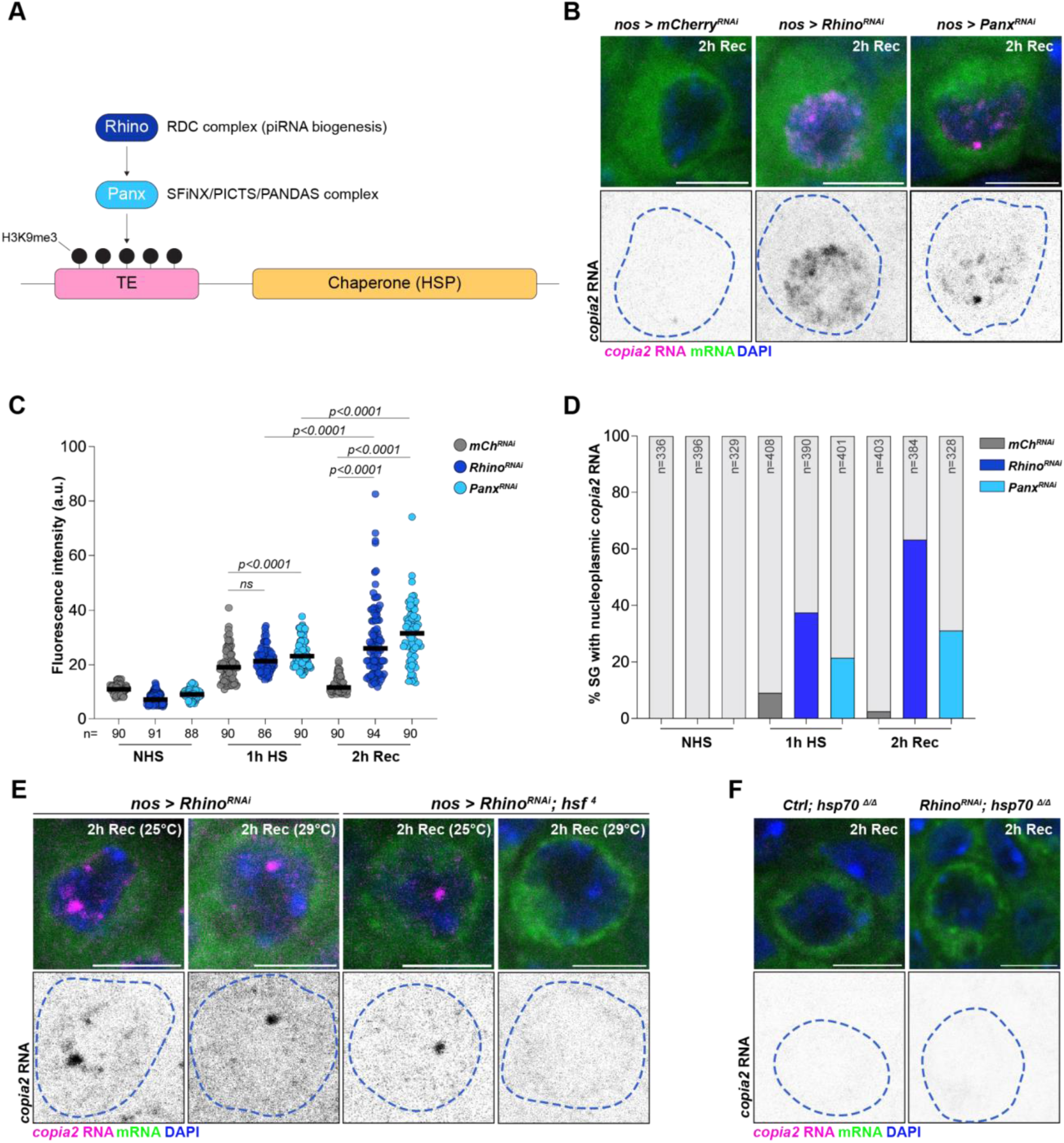
The piRNA pathway ensures silencing of chaperone-embedded TEs during the recovery from heat shock. (A) Schematic of piRNA-dependent deposition of H3K9me3 heterochromatin at TE loci. (B) RNA FISH against *copia2* transcripts (magenta) in spermatogonia from the indicated genotypes at two hours post heat shock. Blue dashed lines demarcate germ cells. Scale bars:5µm. (C, D) Quantification of nucleoplasmic *copia2* expression (C) and percentage of spermatogonia with nucleoplasmic *copia2* expression (D) in the indicated genotypes at the indicated time points. *p-*values were obtained from a two-way ANOVA. ns indicates *p*>0.05. (E) RNA FISH against *copia2* transcripts (magenta) in Rhino-depleted spermatogonia in a *WT* or *hsf* ^4^ background during recovery from heat shock (2h Rec) at either 25°C (permissive temperature) or 29°C (non-permissive temperature). Blue dashed lines demarcate germ cells. Scale bars:5µm. (F) RNA FISH against *copia2* transcripts (magenta) in Control or Rhino-depleted spermatogonia in a *hsp70^Δ/Δ^* background at two hours post heat shock. Blue dashed lines demarcate germ cells. Scale bars:5µm.

We next set out to determine if *copia2* expression during recovery from heat shock in Rhino-depleted germ cells was solely dependent on Hsf. To do so, we depleted Rhino in germ cells in a *hsf* ^4^ background. During recovery from heat shock at 29°C (the non-permissive temperature for *hsf* ^4^), we observed an absence of *copia2* expression in Rhino-depleted germ cells (Fig. 5E), unlike the controls where Hsf function was intact. These data indicated that *copia2* expression during recovery from heat shock, in germ cells lacking a fully functional piRNA pathway, was still dependent on Hsf function. Finally, we asked whether insertions in the *Hsp70B* gene cluster were the main source of *copia2* expression in Rhino-depleted germ cells. To address this, we depleted Rhino in a *hsp70^Δ/Δ^* background, which also lacks the Hsp70-proximal TEs, and observed a complete lack of *copia2* expression during recovery from heat shock (Fig. 5F).

Overall, our data suggest that piRNA pathway-dependent heterochromatin at TE loci blocks TE expression in germ cells during the recovery from heat shock. Moreover, this heterochromatin function, which enables selective chaperone expression in germ cells, requires an nSB-dependent delay in Hsf activity. In germ cells lacking nSBs (e.g. Mitox-treated germ cells), Hsf can override heterochromatin at TE loci during heat shock and drive their stress-induced expression.

### The abundance and accessibility of satellite DNA tune the stress response in germ cells

The formation of nSBs at satellite DNA suggested that these repeats play a crucial role in modulating the germ cell stress response. The majority of the AGAAT repeat, which is bound by Hsf and nucleates nSBs, is present on Chr. 2 and Chr. Y^55,56^ (Fig. 6A). We first tested whether the amount of AGAAT satellite DNA affected nSB formation and function. To do so, we either removed the Y chromosome from males (XO males) or added a Y chromosome to females (XXY females) and assessed stress-induced *Hsp70Bbb* and *copia2* expression in germ cells. Interestingly, spermatogonia from XO males formed fewer nSBs than their XY counterparts while germ cells from XXY females formed more nSBs in comparison to germ cells from XX individuals (Fig. 6B), without significantly affecting nSB size (Fig. S5A). Notably, we observed an inverse correlation between the number of nSBs across these karyotypes and Hsf-dependent and stress-induced expression of *Hsp70Bbb* and *copia2* (Fig. 6C, D, Fig. S5B, C). These data support the idea that AGAAT abundance influences the germ cell-specific transcriptional response to stress but do not rule out other potential Y chromosome contributions.

**Figure 6.**
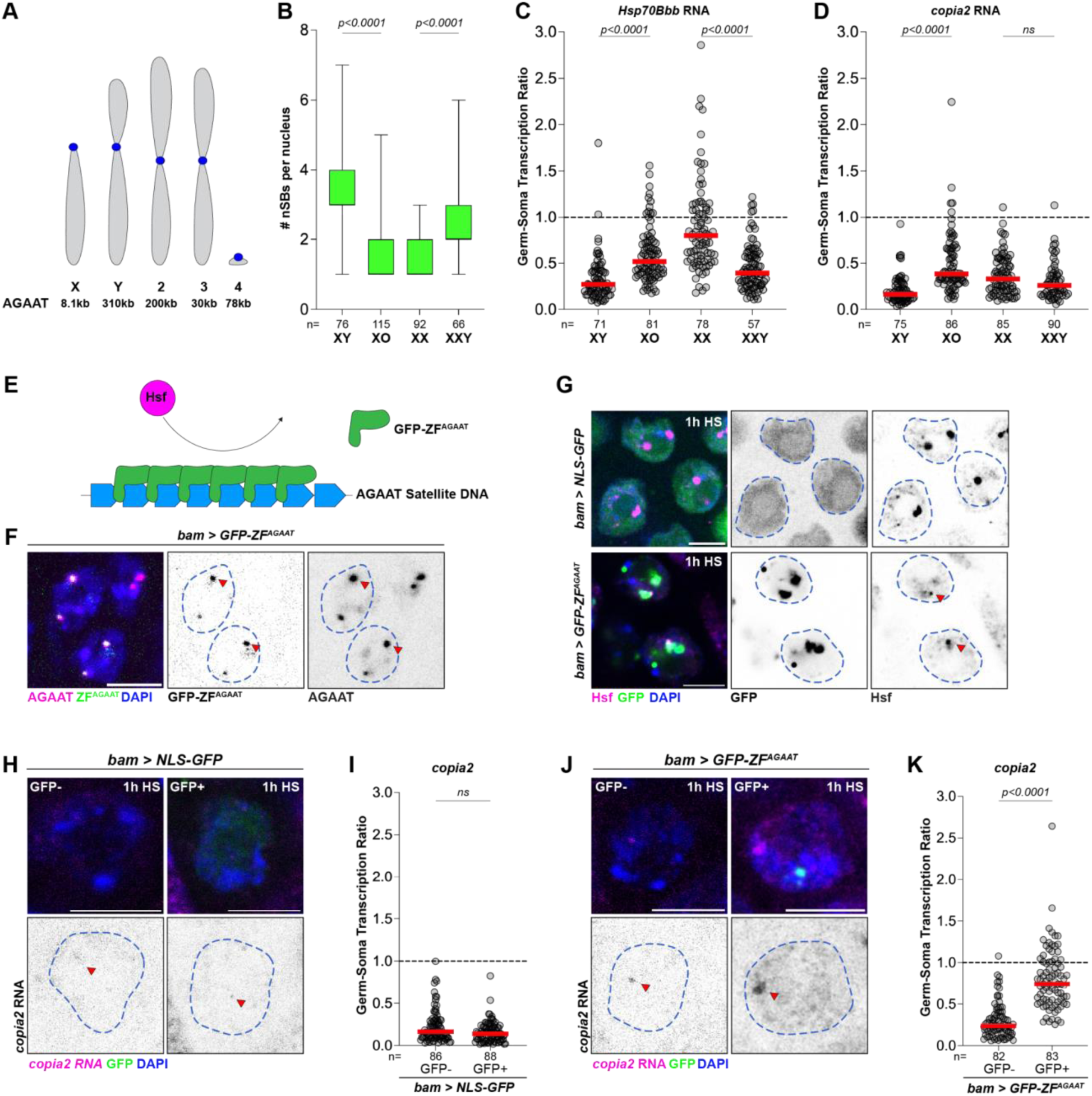
AGAAT satellite DNA abundance and accessibility tunes the stress response in germ cells. (A) Schematic of the *Drosophila melanogaster* karyotype with the abundance of AGAAT (kB) indicated on each chromosome. (B) Quantification of the number of nSBs in male (XO and XY) and female (XX and XXY) germ cells at 1h HS. *p*-values were obtained from a one-way ANOVA. (C, D) Quantification of the germ-soma transcription ratio from nascent *Hsp70Bbb* (C) and *copia2* (D) transcription in males and females from the indicated genotypes. Red lines indicate the median. p-values were obtained from a one-way ANOVA. ns indicates *p*>0.05. (E) Schematic depicting the use of GFP-ZF^AGAAT^ to modulate AGAAT accessibility in germ cells. (F) DNA FISH against the AGAAT satellite DNA in spermatogonia expressing GFP-ZF^AGAAT^ under the control of the *bam-Gal4* driver. Arrowheads show colocalization of GFP-ZF^AGAAT^ and AGAAT satellite DNA. Scale bar: 5µm. (G) Spermatogonia expressing NLS-GFP or GFP-ZF^AGAAT^ (green) stained for Hsf (magenta) and DAPI (blue). Arrowheads indicate impaired nSB formation. Blue dashed lines demarcate germ cells. Scale bars:5µm. (H-K) RNA FISH (H, J) and quantification of germ-soma transcription ratio (I, K) of nascent *copia2* expression (magenta) in spermatogonia expressing NLS-GFP (H, G) or GFP-ZF^AGAAT^ (J, K). GFP-indicates early spermatogonia lacking expression while GFP+ indicates late spermatogonia expressing either NLS-GFP or GFP-ZF^AGAAT^. Arrowheads point to nascent *copia2* transcripts in germ cells, which are demarcated by blue dashed lines. Scale bars:5µm. Red lines indicate the median. *p*-values were obtained using a Welch’s t-test.

To more directly test the role of AGAAT in modulating stress-induced Hsf activity in germ cells, we developed a GFP-tagged Zinc Finger protein (GFP-ZF^AGAAT^, Fig. 6E) to block Hsf access to the AGAAT repeat. Using DNA FISH, we found that GFP-ZF^AGAAT^ localized to the AGAAT satellite DNA (Fig. 6F), but not another similar satellite DNA repeat (Fig. S5D). When expressed in spermatogonia, GFP-ZF^AGAAT^ impaired nSB formation in comparison to the NLS-GFP control (Fig. 6G). Furthermore, spermatogonia expressing GFP-ZF^AGAAT^ exhibit increased nascent expression of Hsp70Bbb (Fig. S5E-H) and copia2 (Fig. 6H-K) in comparison to non-expressing (GFP-) spermatogonia from the same tissue as well as spermatogonia expressing NLS-GFP. Altogether, these experiments establish a crucial role for satellite DNA repeats in tuning the transcriptional response of germ cells to stress via nSB formation.

### Impaired nSB formation is associated with persistent *copia2* expression and cell death

Finally, we set out to understand the functional consequences of nSB perturbation on germ cells following heat shock. First, we quantified nucleoplasmic *copia2* expression in GFP-ZF^AGAAT^-expressing germ cells during the recovery from heat shock (Fig. 7A, B). At 2h post heat shock, we observed that a large fraction of GFP+ (and nSB-deficient) spermatogonia (Fig. 7C) exhibited much higher levels of nucleoplasmic *copia2* transcripts (Fig. 7d). We interpret these results to suggest that when expression of chaperone-proximal TE insertions is triggered in germ cells during heat shock, these TE loci may continue to be expressed during recovery, potentially due to a residual change in the chromatin state. At later stages during recovery from heat shock, we found that testes containing nSB-deficient spermatogonia (GFP-ZF^AGAAT^ or *hsf* ^4^, Fig. 7E, F) exhibited higher levels of germ cell death, as detected by Lysotracker staining^65^. Together, these data highlight the importance of satellite DNA-nucleated nSBs in safeguarding the viability and function of germ cells.

**Figure 7.**
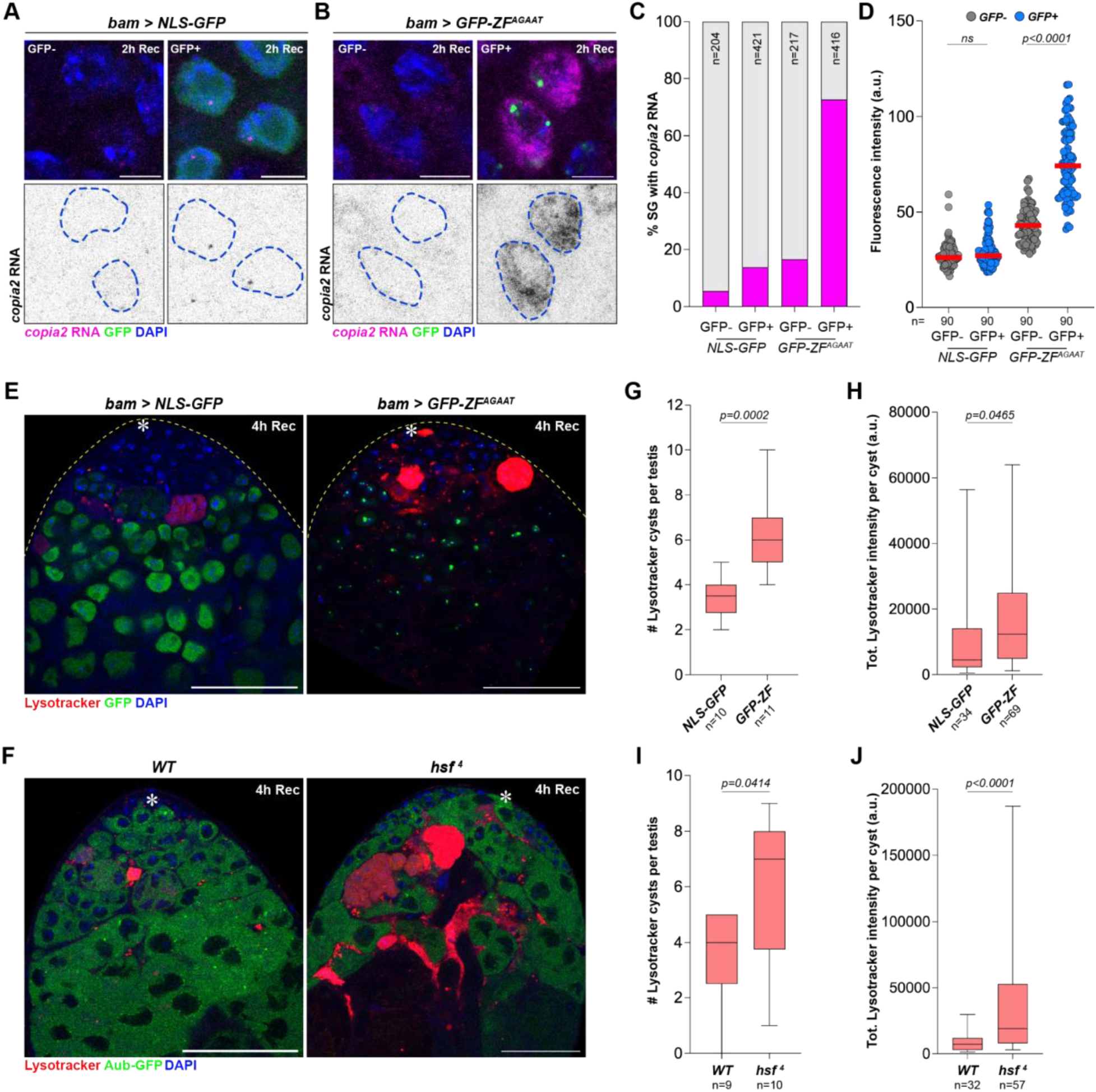
Impaired nSB formation is associated with germ cell death. (A, B) RNA FISH against *copia2* transcripts (magenta) in spermatogonia expressing either NLS-GFP (A) or GFP-ZF^AGAAT^ (B) under the control of the *bam-Gal4* driver at 2h post heat shock. Scale bars:5µm. (C) Quantification of the percentage of spermatogonia with nucleoplasmic *copia2* expression from the indicated genotypes at 2h post heat shock. (D) Quantification of the fluorescence intensity of nucleoplasmic *copia2* expression in spermatogonia from the indicated genotypes at 2h post heat shock. Red lines indicate the median. *p*-values were obtained using a two-way ANOVA. ns indicates *p*<0.05. (E) Lysotracker (red)-stained testes at 4h post heat shock expressing either NLS-GFP or GFP-ZF^AGAAT^ (green) under the control of the *bam-Gal4* driver. Scale bars:25µm. (F) Lysotracker (red)-stained *Aub-GFP* or *hsf* ^4^; *Aub-GFP* testes at 4h post heat shock. Scale bars:25µm. (G-J) Quantification of number of Lysotracker-positive germ cell cysts per testis (G, I) and total Lysotracker intensity per cyst (H, J) from the indicated genotypes at 4h post heat shock. *p*-values were obtained from a Welch’s t-test.

## Discussion

The germline must maintain its function under normal and stress conditions to ensure the safe and faithful passage of genetic information across generations. In this study, we find that *Drosophila* germ cells adopt a distinct transcriptional response to stress, which relies on the formation of Hsf-containing nuclear stress bodies (nSBs) at non-coding satellite DNA repeats. nSBs enable germ cells to selectively express molecular chaperones over stress-induced and chaperone-proximal transposable elements. Notably, germ cells deprived of their ability to form nSBs exhibit elevated levels of cell death upon exposure to stress. While the precise cause for germ cell death remains to be determined, our findings nevertheless emphasize the importance of these germ cell-specific foci in maintaining robust gametogenesis under sub-optimal conditions.

Our data indicate that the primary role of nSBs is to delay Hsf-dependent transcription. Previous reports in human cells have identified that several transcriptional regulators, in addition to Hsf, are enriched at nSBs under stress conditions^42,50,66^. Therefore, we consider it likely that nSBs act as a ‘molecular sink’, sequestering proteins that are required for the transcriptional response to stress in germ cells. Upon post-stress nSB disassembly, the liberation of these transcriptional regulators in germ cells permits Hsf-dependent chaperone expression. Our findings also pinpoint H3K9me3-decorated heterochromatin as a means for Hsf to discriminate between chaperones and TE loci during the recovery from stress. Intriguingly, the transcription inhibitory effects of H3K9me3 appear to be fully functional only at ambient temperatures. At elevated temperatures, Hsf drives the expression of TE loci in both somatic cells and nSB-lacking germ cells. We propose that heat may de-compact heterochromatin, reducing the barrier it poses to the transcription machinery, thus allowing Hsf to stimulate TE expression. Alternatively, the insulation between heterochromatin and euchromatin may be eroded at higher temperatures, allowing euchromatin-bound transcription factors to spread into proximal heterochromatin and drive transcription. Consistently, increasing temperature reduces the spatial compartmentalization of H3K9me3-modified heterochromatin in vitro^67^, while transcription of heterochromatin loci upon stress has been observed in several contexts^20,48,68,69^.

We find the high density of TE insertions within the *Hsp70* gene clusters to be particularly noteworthy and speculate that this phenomenon occurs due to the increased chromatin accessibility of these regions upon stress-dependent transcription^70^. Once successfully embedded in proximity to a chaperone, TEs can couple their expression to the canonical Hsf-dependent stress response, thereby facilitating their propagation in the host genome. While the reference *Drosophila* genome mainly contains *copia2*, *S-element* and *invader1* insertions in the *Hsp70* gene clusters^31^, natural populations harbor different TEs at the same loci^71–75^, likely due to opportunistic insertions following exposure to environmental stress in their native habitats. In these cases, nSBs may function as a prophylactic genome defence mechanism, blocking excessive TE expression and mobilization upon further stress exposure. This function may be particularly significant since the piRNA pathway, which typically protects the germline genome from transposon mobilization, has been suggested to be partially impaired at elevated temperatures^17,27^.

Finally, our study suggests a potentially broad role for satellite DNA repeats in tuning cellular processes, based on their ability to mimic transcription factor binding sites. The formation of nuclear stress bodies, both in primate somatic cells and in the *Drosophila* germline, relies on the similarity between NGAAN satellite DNA repeats and the Hsf binding motif. Similarly, other transcription factors localize to satellite DNA repeats based on repeated binding motifs and this sequestration influences cellular function^76,77^. What’s perhaps most intriguing about these satellite DNA-dependent mechanisms is that the underlying repeats evolve rapidly, such that their ability to recruit transcription factors or other DNA-binding proteins may vary substantially between individuals. This raises the possibility that differences in satellite DNA composition may contribute to phenotypic differences between individuals that cannot be otherwise explained by changes in protein-coding sequences. Consistent with this idea, introgression of the AGAAT-rich but gene-poor *Drosophila* Y chromosome from tropical, but not temperate, populations into an otherwise identical genetic background enhances fertility at higher temperatures^78^. Beyond explaining phenotypic variation between individuals, we also consider that rapid changes in satellite DNA copy number may allow individuals and populations to adapt to their environment on short timescales, thus maintaining evolutionary fitness in the face of an ever-changing environment.

## Methods

### Fly husbandry and strains

All fly strains were raised on standard Bloomington medium unless otherwise indicated. All crosses were conducted at 25°C and 70% humidity. For heat shock, 0-3 day old adult flies and third instar larvae were subjected to a heat shock by placing them in a water bath set to 37°C for one hour. The vials were shifted to an incubator set to 25°C for recovery. For embryo heat shock, flies were allowed to lay eggs on standard apple juice agar plates with fresh yeast paste for approximately 16 hours. Adult flies were then removed and the plates containing embryos were subjected to a one hour heat shock in an incubator set to 37°C, followed by recovery in an incubator set to 25°C. *Oregon R* was used as a wild type strain. *bam^Δ^*^86^ (BDSC5427), *c587-Gal4* (BDSC67747), *UASt-GFP-Rpl10Ab* (BDSC42683), *hsf* ^4^ (BDSC5490), *hsp70^Δ^ (Df(3R)Hsp70A, Df(3R)Hsp70B*, BDSC8841), *UAS-mCD4-tdTomato* (BDSC98368), *UAS-mCherry^RNAi^* (BDSC35785), C(1)RM (BDSC9460), *UASt-NLS-GFP* (BDSC4776) were obtained from the Bloomington Drosophila Stock Center. Aubergine-GFP (VDRC313243) UAS-*Rhino^RNAi^* (VDRC330007) UAS-*panoramix^RNAi^* (VDRC313142) were obtained from the Vienna Drosophila Research Center. *nos-Gal4* (Chr. 2), *nos-Gal4* (Chr. 3), and *bam-Gal4* were gifts from Yukiko Yamashita. *bam*^1^ has been previously described^79^.

### Transgene construction

For *UASt-GFP-Hsf*, the Hsf PB isoform (Q4H2F9) was synthesized (Twist Biosciences) and subcloned into the *pUASt-EGFP-attB* vector^80^ downstream of the BglII site. Transgenic flies were generated by PhiC31 integrase-mediated transgenesis into the attP40 site (Bestgene). For *UASt-GFP-ZF^AGAAT^*, we used previously described Zinc Finger design principles^81^. We used a custom python script to determine which register of the AGAAT repeat can be directly targeted using the DNA-binding motifs of established ZFs. We used an 18bp repeat sequence, which previously was shown to enable successful targeting^77^. We then used AlphaFold to predict binding strength between five ZF candidates and their respective 18bp target sequences. The ZF array targeting 5’ AAT AGA ATA GAA TAG AAT 3’ had the highest pTM and ipTM scores, and was chosen for experimental validation. The resulting amino acid sequence was codon optimized for Drosophila, synthesized (Twist Biosciences) and subcloned into the *pUASt-EGFP-attB* vector^80^ downstream of the BglII site. Transgenic flies were generated by PhiC31 integrase-mediated transgenesis into the attP2 site (Bestgene).

### RNA extraction, sequencing and analysis

50 to 60 testes from 0-3 day old males were dissected into RNAse-free 1x PBS and flash frozen in liquid nitrogen. RNA was extracted from the testes using the Qiagen RNeasy RNA extraction kit. The samples were processed using the Promega ReliaPrep RNA Clean Up and Concentration System Kit. RNA concentrations were determined using both Nanodrop and Qubit. Samples of sufficient quality (RIN value > 9) were used to perform library preparation (Illumina TruSeq mRNA kit), which was followed by sequencing using Illumina NovaSeq 6000 (paired end, 250bp) at the Functional Genomics Center Zurich (FGCZ). The resulting raw reads were processed by removing adaptor sequences, low-quality-end trimming, and removal of low-quality reads using BBTools v38.18. The exact commands used for quality control can be found on the Methods in Microbiomics webpage^82^. The quality-controlled reads were aligned against BDGP6.32 using STAR aligner v. 2.7.8 aligner^83^. Transcript and Transposon abundances were quantified using TEtranscripts v. 2.2.3^33^. Differential gene analysis was performed using the Bioconductor R package DESeq2 v1.37.4^84^. Simple satellite RNA transcripts (k-mers) were identified from the RNAseq dataset using the k-seek pipeline^85^.

### RNA *in situ* hybridization

RNA *in situ* hybridization in testes and ovaries was performed as previously described^86^ with minor modifications. Briefly, testes/ovaries were dissected in 1xPBS, and treated with 4% formaldehyde prepared in RNase-free 1xPBS. Post fixation, samples were washed twice with 1xPBS for 5 mins each before adding 70% ethanol diluted in RNase-free water and nutating overnight at 4°C. The following day, the samples were washed with RNA FISH wash buffer (2xSSC, 10% formamide) for 5 minutes at room temperature. The samples were then incubated in hybridization buffer (50nM probes, 2xSSC, 10% dextran sulfate, 1g/L *S.cerevisiae* tRNA, 2mM Vanadyl ribonucleoside complex, 0.5% RNase-free Ultrapure BSA, and 10% deionized formamide) at 37°C for 12-16 hours. Next, the samples were washed twice with RNA FISH wash buffer at 37°C for 30 mins each and then mounted in VECTASHIELD with DAPI (Vector Labs). For smRNA FISH, the following probe was used to label the ATTCT RNA (5’ Cy5-AAT AGA ATA GAA TAG AAT AGA ATA GAA TAG-Cy5 3’). A fluorophore-coupled oligo-dT probe was used to label polyadenylated RNA (mRNA). Stellaris RNA probes (LGC Biosearch Technologies) were designed against the coding sequence of *Hsp70Bbb* (NM_176486) and the consensus sequence of *copia2*^87^. Samples were imaged using a Leica SP8 confocal microscope with a 63x oil-immersion objective (NA=1.4).

### Immunofluorescence staining

0-3 day old males, 3-4 day old females or wandering 3^rd^ instar larvae were used for dissection of testes, ovaries and somatic tissue respectively. Briefly, the samples were dissected in 1x PBS, transferred to 4% EM-grade paraformaldehyde in 1xPBS and incubated with nutating for 30 minutes. The fixed samples were then washed three times for 20 minutes each in 1x PBS-T (PBS containing 0.1% Triton-X 100). The samples were then incubated in 3% Bovine Serum Albumin in 1x PBS-T for 60 minutes for blocking. Primary antibodies diluted in 3% Bovine Serum Albumin in 1x PBS-T were then added to the samples and incubated at 4°C overnight. The next day, samples were washed three times for 15 minutes each with 1x PBS-T and incubated overnight at 4°C with secondary antibodies diluted in 3% BSA in 1X PBS-T. Samples were washed again and stored in DAPI-containing mounting medium. Fixation of *Drosophila* embryos was performed according to standard protocol. Briefly, collected embryos were moved to glass scintillation vials, washed with Embryo Wash Buffer (7% NaCl, 0.5% Triton-X 100), and then dechorionated by vigorously shaking in 3 ml 50% bleach solution for 90 seconds. Embryos were then washed five times in 1x PBS to remove bleach. Dechorionated embryos were fixed in a 1:1 solution of heptane:4% EM-grade paraformaldehyde by vigorously shaking for 20 minutes. The bottom aqueous layer was removed post fixation, and ice-cold methanol was added to devitellinize the embryos. After this, embryos were washed 3 times in ice-cold methanol and stored in methanol at -20°C until further use. Embryos were rehydrated using serial washes of methanol:1x PBST-T at 4:1, 1:1 and 1:4 volumes, followed by one wash in 1x PBS-T for 5 minutes. Embryos were then subjected to immunofluorescence staining as described above. For lysotracker staining, testes were dissected in 1x PBS and incubated in Lysotracker for 30 minutes at room temperature. Samples were then rinsed with 1x PBS twice, followed by fixation and staining as described above. The following antibodies were used in this study: Rabbit anti-Hsf (1:500, gift from Carl Wu) and Rat anti-Vasa (1:100, AB_760351, DSHB).

### Image analysis and quantification

All image analysis was performed in FIJI. Quantification of the Germ-Soma Transcription Ratio was performed as follows. For each sample, z-stacks were acquired for the entire apical tip (∼40-50 slices, 0.5µm thickness). We then selected a somatic cell and a germ cell that were maximally 3 slices apart to avoid variability in fluorescence intensity arising from distance from the objective. We demarcated a region of interest (ROI) around the nascent locus of transcription in the selected somatic and germ cell. The Germ-Soma Transcription Ratio was calculated by dividing the total fluorescence intensity of the germ cell nascent locus by that of the somatic cell nascent locus for each pair. Around 60 such pairs were quantified from 5 to 6 testes. For quantification of *Hsp70Bbb* cytoplasmic expression, the average fluorescence intensity was measured for an ROI of 0.2-0.4µm^2^ in the cytoplasm of somatic or germ cells. Nucleoplasmic *copia2* RNA expression was quantified by generating an ROI around the entire nucleus of a somatic or germ cell and measuring the average fluorescence intensity. nSB size was measured using the particle analysis function in FIJI. Briefly, a fluorescence intensity threshold was set for each z-stack of the apical tip, to identify nSBs as particles, after which the area of each nSB per nucleus was measured. The area of each nSB was then normalized by the area of its respective nucleus at its maximal diameter. Somatic expression of *UAS-mCD4-tdTomato* under the control of *c587-Gal4* was used to stage spermatogonial cysts and quantify normalized nSB size and number. The vast majority of images are from a single plane while the remainder are a maximum intensity projection of 2-3 slices at most. All statistical tests were conducted using Graphpad Prism.

### Mitoxantrone treatment and heat shock

Testes from 5-6 males were dissected in 1x PBS and transferred to vials containing 1ml Schneider’s insect medium supplemented with either 20mM Mitoxantrone or ddH_2_O (Control). Vials were incubated on a nutator at room temperature for 90 minutes and then transferred to a dry bath at 37°C for one hour. No Heat Shock (NHS) samples were incubated at 25°C for the same amount of time. After heat shock, the Schneider’s insect medium was removed, samples were rinsed twice with 1x PBS and subjected to the previously described RNA *in situ* hybridization or immunofluorescence staining protocols.

### Satellite DNA FISH

10-14 testes were dissected and fixed as described above, and optional immunofluorescence staining protocol was carried out first. Subsequently, samples were fixed with 4% formaldehyde for 10 minutes and washed in 1xPBS-T for 30 minutes. Fixed samples were incubated with 2 mg/ml RNase A solution at 37°C for 10 minutes, then washed with 1x PBS-T +1 mM EDTA. Samples were washed in 2xSSC-T (2xSSC containing 0.1% Tween-20) with increasing formamide concentrations (20%, 40% and 50%) for 15 minutes each followed by a final 30 minute wash in 50% formamide. Hybridization buffer (50% formamide, 10% dextran sulfate, 2x SSC, 1 mM EDTA, 1 μM probe) was added to washed samples. Samples were denatured at 91°C for 2 minutes, then incubated overnight at 37°C. The following fluorescent probes were used for hybridization: AATAG – 5’ Cy5-AAT AGA ATA GAA TAG AAT AGA ATA GAA TAG 3’ and AATAT – 5’ Cy5-AAT ATA ATA TAA TAT AAT ATA ATA TAA TAT 3’. Following hybridization, samples were washed with 2x SSC-T three times for 15 minutes each. Samples were then mounted and imaged as described before.

## Acknowledgements

We thank members of the Jagannathan lab, Hugo Stocker, Yves Barral for discussion and comments on the manuscript. We are grateful for the reagents and resources provided by Yukiko Yamashita, Carl Wu, the Bloomington Drosophila Stock Center, the Vienna Drosophila Resource Center and the Developmental Studies Hybridoma Bank. We acknowledge microscopy support from the Scientific Center for optical and Electron Microscopy (ScopeM), NGS support from the Functional Genomics Center Zurich (FGCZ) and bioinformatic infrastructure from Shinichi Sunagawa. MJ acknowledges the following funding sources: Swiss National Science Foundation (310030_189131 and 320030_228043).

## Notes

### Competing Interest Statement

The authors have declared no competing interest.

