## Supplementary Figures for "Nuclear stress bodies enable a germline-specific transcriptional stress response in *Drosophila*"

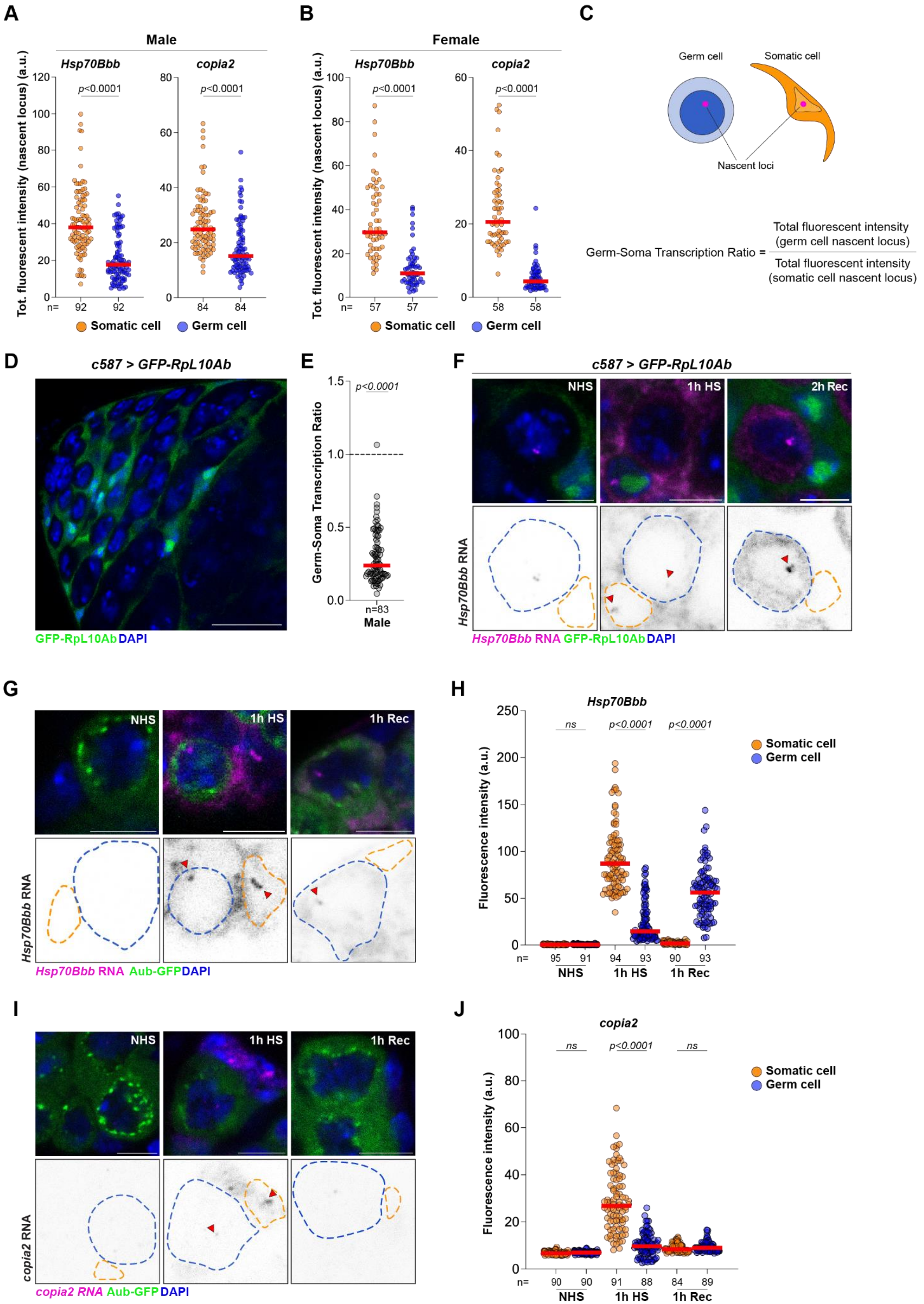

A
Male
Hsp70Bbb
copia2
Tot. fluorescent intensity (nascent locus) (a.u.)
p<0.0001
p<0.0001
n= 92 92 84 84
Somatic cell
Germ cell
B
Female
Hsp70Bbb
copia2
p<0.0001
p<0.0001
n= 57 57 58 58
C
Germ cell
Somatic cell
Nascent loci
Germ-Soma Transcription Ratio = Total fluorescent intensity (germ cell nascent locus) / Total fluorescent intensity (somatic cell nascent locus)
D
c587 > GFP-RpL10Ab
GFP-RpL10Ab DAPI
E
Germ-Soma Transcription Ratio
p<0.0001
n=83
Male
F
c587 > GFP-RpL10Ab
NHS
1h HS
2h Rec
Hsp70Bbb RNA
Hsp70Bbb RNA GFP-RpL10Ab DAPI
G
NHS
1h HS
1h Rec
Hsp70Bbb RNA
Hsp70Bbb RNA Aub-GFP DAPI
H
Hsp70Bbb
Fluorescence intensity (a.u.)
ns
p<0.0001
p<0.0001
Somatic cell
Germ cell
n= 95 91 94 93 90 93
NHS
1h HS
1h Rec
I
NHS
1h HS
1h Rec
copia2 RNA
copia2 RNA Aub-GFP DAPI
J
copia2
Fluorescence intensity (a.u.)
ns
p<0.0001
ns
Somatic cell
Germ cell
n= 90 90 91 88 84 89
NHS
1h HS
1h Rec

**Supplementary Figure 1. Characterization of the stress response in *Drosophila* male and female germ cells**

(A, B) Total fluorescent intensities of nascent transcriptional loci of *Hsp70Bbb* and *copia2* in mitotically proliferating male (A) and female (B) germ cells. These values were used to calculate the germ-soma transcriptional ratio for neighbouring germ and somatic cells. *p*-values were obtained from a Welch's t-test.

(C) Schematic depicting the calculation of the germ-soma transcription ratio.

(D) Expression of GFP-RpL10Ab in somatic cells of the testes using the *c587-Gal4* driver. GFP marks the cytoplasm of somatic cells. Scale bar:25μm

(E) Quantification of the germ-soma transcription ratio for *Hsp70Bbb* in the *c587-Gal4*>GFP-RpL10Ab background at 1h HS. Red line indicates the median. *p*-values were obtained from a one sample *t*-test against a hypothetical value of 1.

(F) RNA FISH against *Hsp70Bbb* transcripts (magenta) at the indicated time point in testes expressing RpL10Ab (green) in somatic cells and stained with DAPI (blue). Blue dashed lines demarcate germ cells while orange dashed lines demarcate somatic cell nuclei.Scale bar:5μm.

(G-J) RNA FISH (G, I) and quantification (H, J) of *Hsp70Bbb* (G, H) and *copia2* (I, J) transcripts at the indicated time points in ovaries expressing Aub-GFP (green) and stained with DAPI (blue). Blue dashed lines demarcate germ cells while orange dashed lines demarcate somatic cell nuclei. Arrowheads indicate nascent transcriptional loci. Scale bar:5μm. *p*-values were obtained from a two-way ANOVA.

**A**

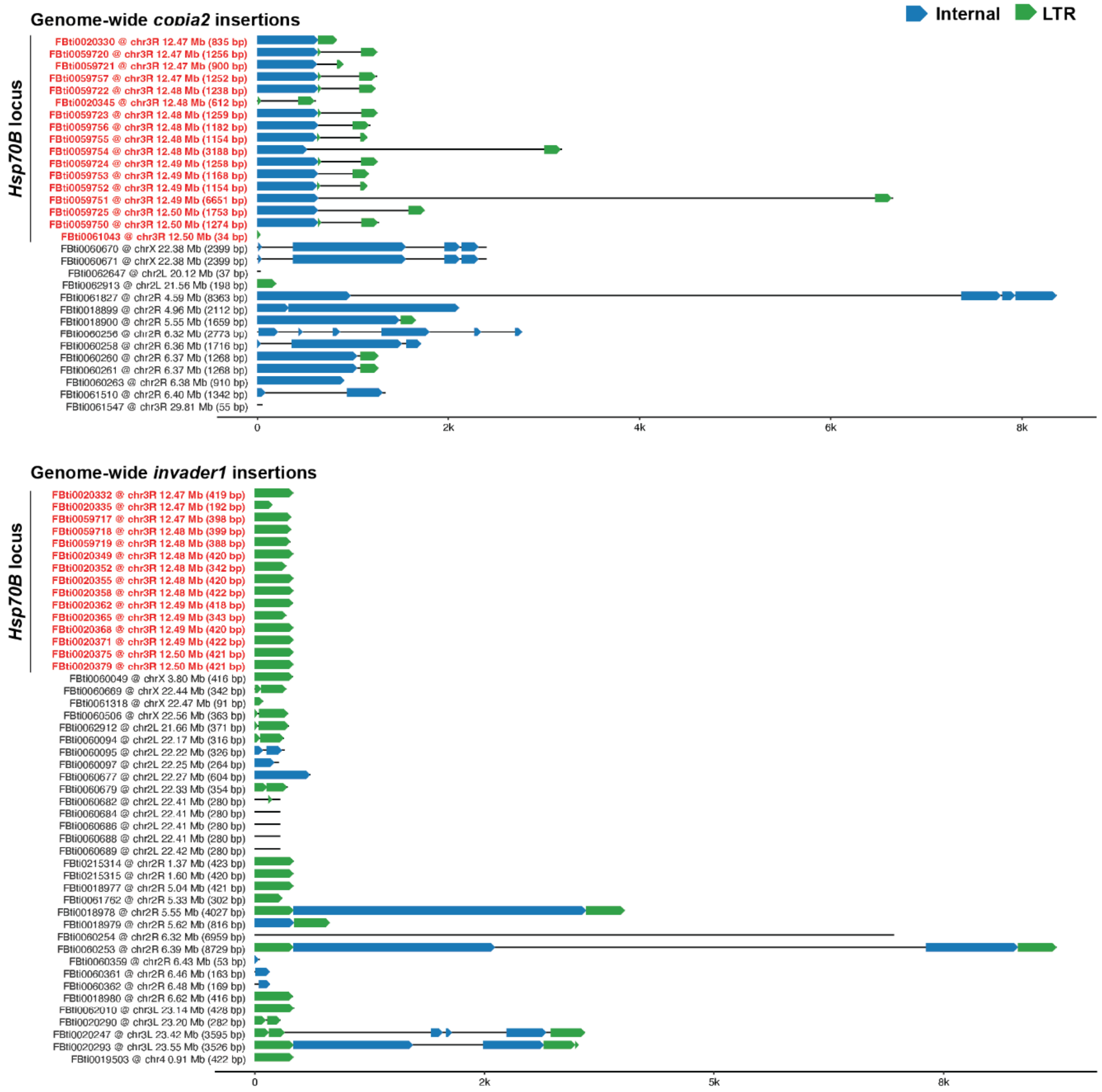


**B**

| Fly Base ID | Symbol | Chromosomal location | $log_2$FoldChange | $p_{adj}$ |
|---|---|---|---|---|
| FBti0059720 | Dmel\Dm88{}1659 | 3R (87B12) | 5.189812171 | 8.57E-53 |
| FBti0059721 | Dmel\Dm88{}1660 | 3R (87B13) | 6.108406994 | 1.91E-29 |
| FBti0020330 | Dmel\Dm88{}1295 | 3R (87B12) | 4.895073315 | 1.09E-13 |
| FBti0019359 | Dmel\S{}1293 | 3R (87A2-87A3) | 8.821087541 | 9.86E-11 |
| FBti0020235 | Dmel\R1A1{}1124 | 3L (80D2) | 2.505946564 | 1.49E-08 |
| FBti0061120 | Dmel\INE-1{}3059 | 2R (43C3) | 8.941504242 | 2.50E-08 |
| FBti0019362 | Dmel\S{}1347 | 3R (87B14) | 3.127949573 | 1.81E-06 |
| FBti0062822 | Dmel\S{}4761 | 3R (87B14) | 6.828559503 | 5.14E-06 |
| FBti0059750 | Dmel\Dm88{}1689 | 3R (87B14) | 2.776907081 | 0.01547995 |
| FBti0059789 | Dmel\INE-1{}1728 | 3R (84B2) | 6.048823613 | 0.03130254 |
| FBti0061113 | Dmel\INE-1{}3052 | 2R (43C3) | 6.234193597 | 0.04039616 |

**Supplementary Figure 2. Genomic coordinates and stress-induced expression of individual TE insertions.**

(A) Genomic coordinates and structure of *copia2* and *invader1* insertions in the *Drosophila melanogaster* genome. Loci marked in red are embedded within the *Hsp70B* gene cluster.

(B) Identity of the 11 TE insertions that were uniquely identified and upregulated following heat shock from our RNA-seq data.

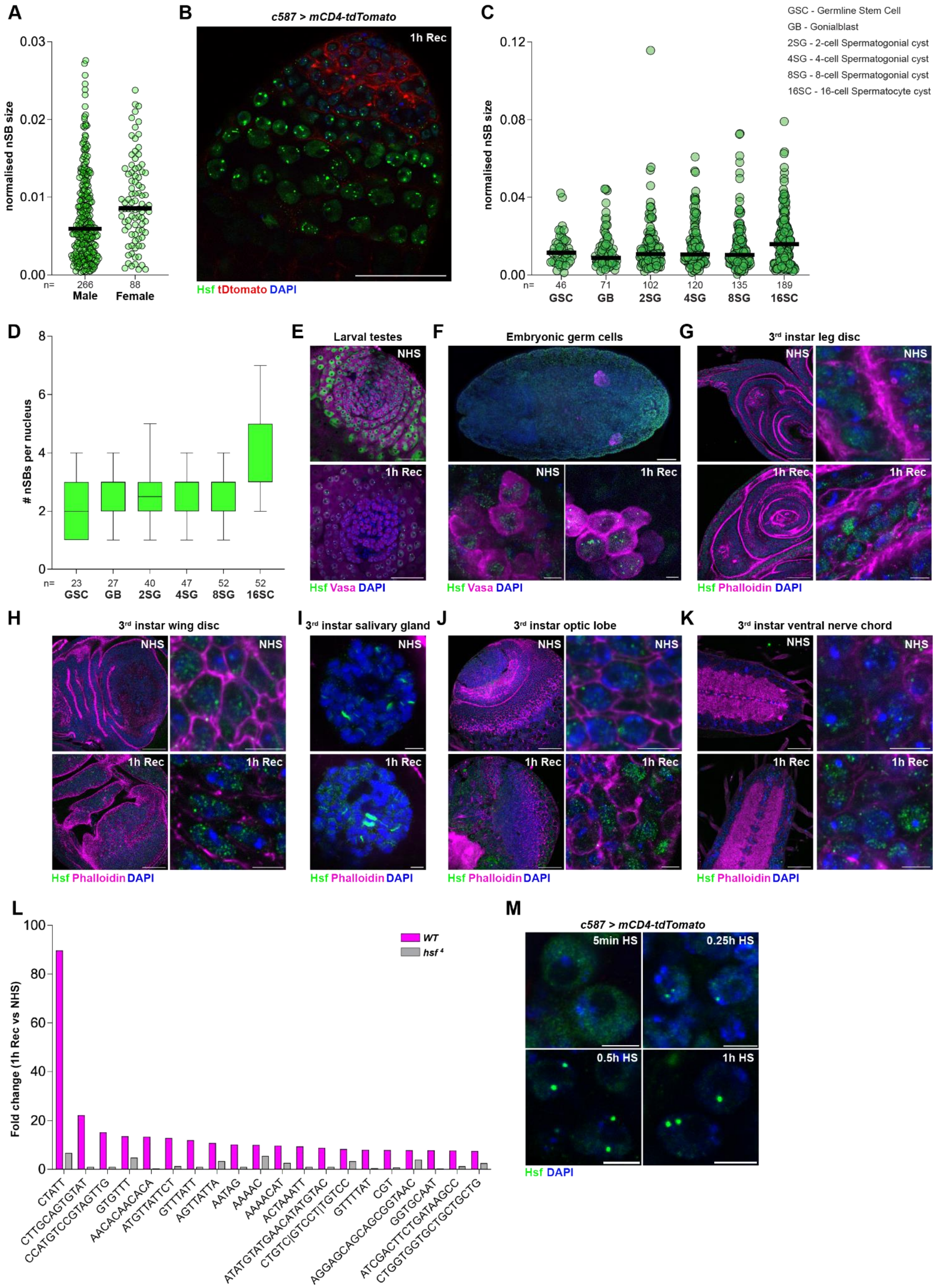

A
normalised nSB size
0.03
0.02
0.01
0.00
n=
266
88
Male
Female
B
c587 > mCD4-tdTomato
1h Rec
Hsf tDtomato DAPI
C
normalised nSB size
0.12
0.08
0.04
0.00
46
71
102
120
135
189
GSC
GB
2SG
4SG
8SG
16SC
GSC - Germline Stem Cell
GB - Gonialblast
2SG - 2-cell Spermatogonial cyst
4SG - 4-cell Spermatogonial cyst
8SG - 8-cell Spermatogonial cyst
16SC - 16-cell Spermatocyte cyst
D
### nSBs per nucleus
23
27
40
47
52
52
E
Larval testes
NHS
1h Rec
Hsf Vasa DAPI
F
Embryonic germ cells
G
3rd instar leg disc
Hsf Phalloidin DAPI
H
3rd instar wing disc
I
3rd instar salivary gland
J
3rd instar optic lobe
K
3rd instar ventral nerve chord
L
Fold change (1h Rec vs NHS)
WT
hsf 4
CTATT
CTTGCAGTGTAT
CCATGTCCGTAGTTG
GTGTT
AACACAACACA
ATGTTATTCT
GTTTATT
AGTTATTA
AATAG
AAAAC
AAAACAT
ACTAAATT
ATATGTATGAACATATGTAC
CTGTC|GTCCT|TGTCC
GTTTTAT
CGT
AGGAGCAGCAGCGGTAAC
GGTGCAAT
ATCGACTTCTGATAAGCC
CTGGTGGTGCTGCTGCTG
M
c587 > mCD4-tdTomato
5min HS
0.25h HS
0.5h HS
1h HS
Hsf DAPI

**Supplementary Figure 3. Hsf-containing nuclear stress bodies are specifically formed in germ cells.**

(A) Quantification of the area of Hsf foci normalized to nuclear area in male and female mitotically proliferating germ cells from an Aub-GFP background.

(B) The apical tip of a testis expressing *mCD4-tdTomato* under the control of the *c587-Gal4* driver, outlining individual germ cell cysts. Scale bar:25μm

(C,D) Quantification of size (C) and number per nucleus (D) of Hsf foci in the indicated germ cell stages in the *c587-Gal4* > *mCD4-tdTomato* background.

(E, F) Larval testes (E) and Stg.15-16 embryos (F) from Oregon R stained for Hsf (green), Vasa (magenta) and DAPI (blue) at the indicated time points. Scale bars:5μm

(G-K) The indicated tissues stained for Hsf (green), Vasa (magenta) and DAPI (blue) before (NHS) and 1h post heat shock (1h Rec). Scale bars:5μm

(L) Average fold change of k-mer abundance from the RNA-seq data from Fig. 1B, C. k-mer abundance was normalized by the total number of uniquely mapped reads for each replicate.

(M) Spermatogonia stained for Hsf (green) and DAPI (blue) at the indicated time points in the *c587-Gal4* > *mCD4-tdTomato* background.

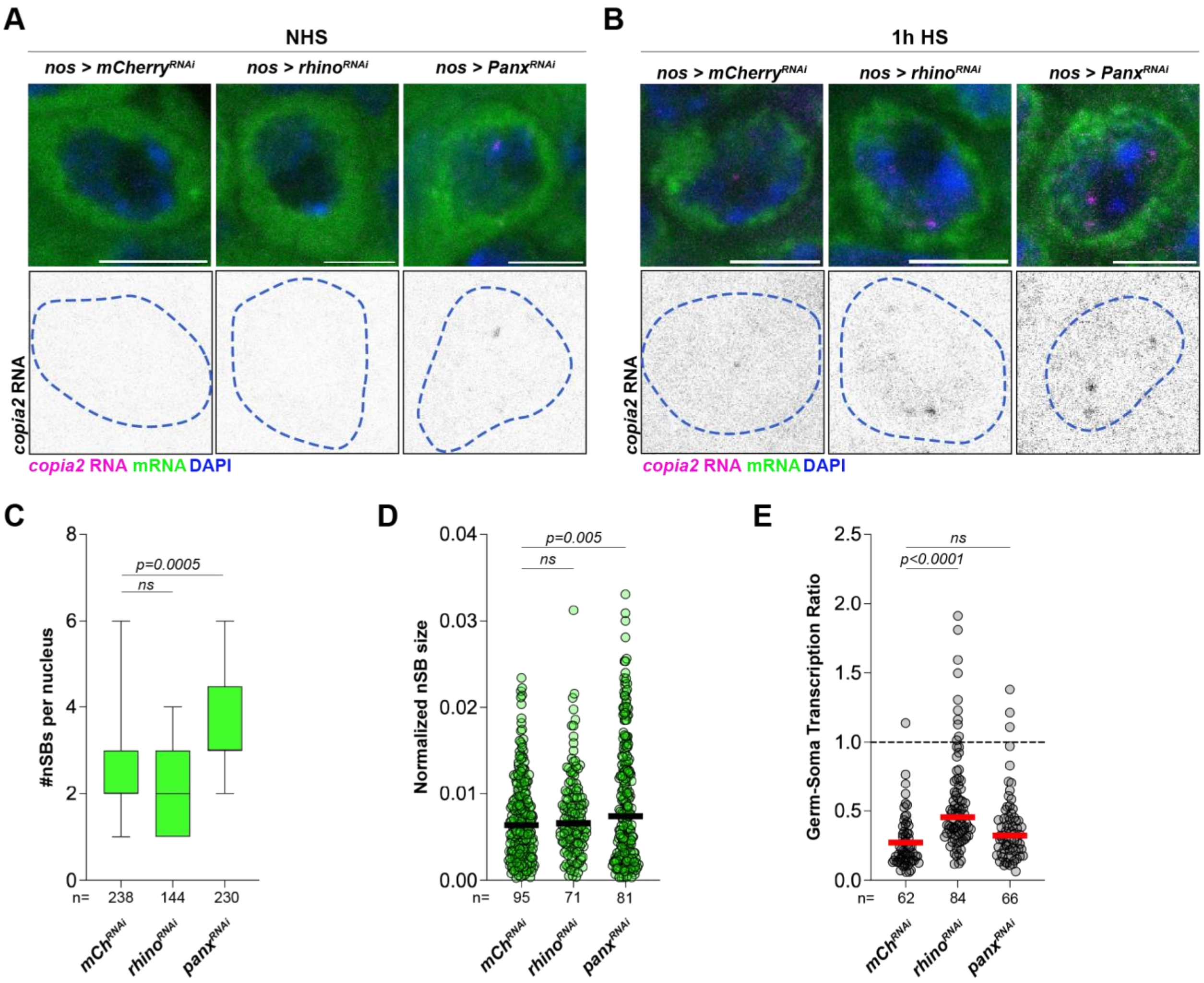


**Supplementary Figure 4. Characterization of *copia2* expression and nSB formation in germ cells lacking piRNA pathway components.**

(A, B) RNA FISH against *copia2* transcripts (magenta) in spermatogonia from the indicated genotypes in the absence of heat shock (A, NHS) and immediately after heat shock (B, 1h HS). Blue dashed lines demarcate germ cells. Scale bars:5µm.

(C) Quantification of the number of nSBs per spermatogonial nucleus expressing the indicated RNAi under the control of the *nos-Gal4* driver. *p*-values were obtained from a one-way ANOVA. ns indicates *p*<0.05.  in the indicated genotypes.

(D) Quantification of nSB size normalized to nuclear area in spermatogonia expressing the indicated RNAi under the control of the *nos-Gal4* driver. Black lines indicate median values. *p*-values obtained using a one-way ANOVA.

(E) Quantification of the germ-soma transcription ratio in spermatogonia expressing the indicated RNAi under the control of the *nos-Gal4* driver. Red lines indicate median values. *p*-values obtained  using a one-way ANOVA.

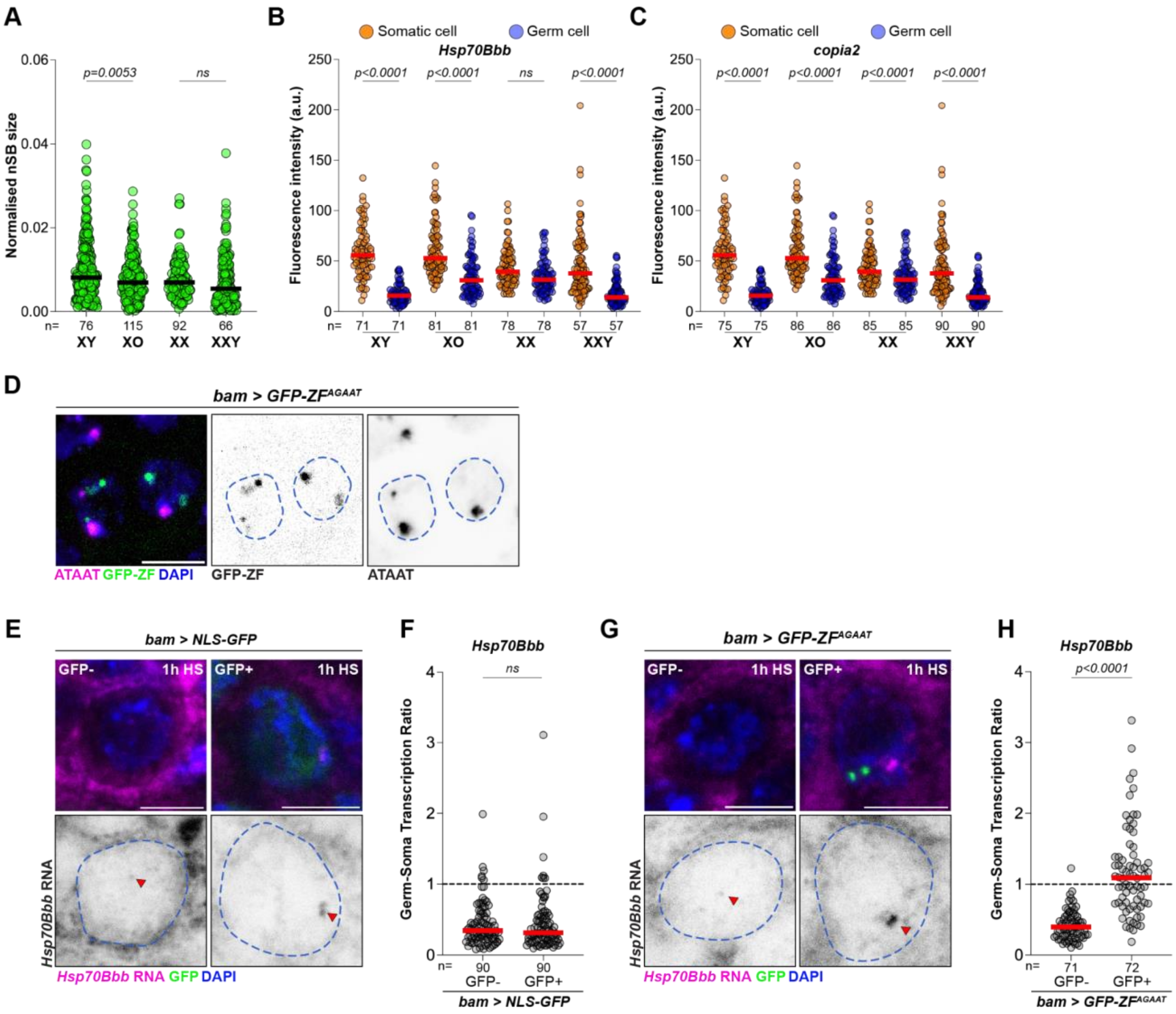


**Supplementary Figure 5. nSB formation and function is regulated by the AGAAT satellite DNA repeat.**

(A) Quantification of normalised nSB size in male (XO and XY) and female (XX and XXY) germ cells immediately after heat shock. Black lines indicate the median. *p*-values were obtained using a one-way ANOVA. ns indicates $p>0.05$.

(B,C) Fluorescence intensity of nascent transcriptional loci of *Hsp70Bbb* (B) or *copia2* (C) in somatic cells (orange) and germ cells (blue) from the indicated genotypes at 1h HS. These values were used to calculate the germ-soma transcriptional ratio for neighbouring germ and somatic cells. Red lines indicate the median. *p*-values were calculated using a two-way ANOVA. ns indicates $p>0.05$.

(D) DNA FISH against the ATAAT satellite DNA in spermatogonia expressing GFP-ZF$^{AGAAT}$ under the control of the *bam-Gal4* driver. Scale bar: 5µm.

(E-G) RNA FISH (E, G) and quantification of germ-soma transcription ratio (F, G) of nascent *Hsp70Bbb* expression (magenta) in spermatogonia expressing NLS-GFP (E, F) or GFP-ZF$^{AGAAT}$ (G, G). GFP-indicates early spermatogonia lacking expression while GFP+ indicates late spermatogonia expressing either NLS-GFP or GFP-ZF$^{AGAAT}$. Arrowheads point to nascent *Hsp70Bbb* transcripts in germ cells, which are demarcated by blue dashed lines. Scale bars:5µm. Red lines indicate the median. *p*-values were obtained using a Welch's t-test.
